# A continuous centennial Lateglacial-Early Holocene (15-10 cal kyr BP) palynological record from the Iberian Pyrenees and regional comparisons

**DOI:** 10.1101/2023.07.02.547433

**Authors:** Valentí Rull, Arnau Blasco, Miguel Ángel Calero, Maarten Blaauw, Vegas-Vilarrúbia Teresa

## Abstract

This paper presents the first continuous (gap-free) Lateglacial-Early Holocene (LGEH) pollen record for the Iberian Pyrenees resolved at centennial resolution. The main aims are (i) to provide a standard chronostratigraphic correlation framework, (ii) to unravel the relationships between vegetation shifts, climatic changes and fire, and (iii) to obtain a regional picture of LGEH vegetation for the Pyrenees and the surrounding lowlands. Seven pollen assemblage zones were obtained and correlated with the stadial/interstadial phases of the Greenland ice cores that serve as a global reference. Several well-dated datums were also derived for keystone individual taxa that are useful for correlation purposes. Four vegetation types were identified, two of them corresponding to conifer and deciduous forests (Cf, Df) and two representing open vegetation types (O1, O2) with no modern analogs, dominated by *Artemisia*-Poaceae and *Saxifraga*-Cichiroideae, respectively. Forests dominated during interstadial phases (Bølling/Allerød and Early Holocene), whereas O1 dominated during stadials (Oldest Dryas and Younger Dryas), with O2 being important only in the first half of the Younger Dryas stadial. The use of pollen-independent proxies for temperature and moisture allowed the reconstruction of paleoclimatic trends and the responses of the four vegetation types defined. The most relevant observation in this sense was the finding of wet climates during the Younger Dryas, which challenges the traditional view of arid conditions for this phase on the basis of former pollen records. Fire incidence was low until the early Holocene, when regional fires were exacerbated, probably due to the combination of higher temperatures and forest biomass accumulation. These results are compared with the pollen records available for the whole Pyrenean range and the surrounding lowlands within the framework of elevational, climatic and biogeographical gradients. Some potential future developments are suggested on the basis of the obtained results, with an emphasis on the reconsideration of the LGEH spatiotemporal moisture patterns and the comparison of the Pyrenees with other European ranges from different climatic and biogeographical regions.

## 1. Introduction

In the Iberian Peninsula (IP), the Lateglacial, or the Last Glacial Maximum (LGM)-Holocene transition, is an excellent time interval to study the response of vegetation to natural drivers of ecological change due to the occurrence of conspicuous climatic reversals and the lack of significant human pressure [1]. This is especially true for mountain ranges, where extensive landscape anthropization did not begin until the Late Holocene [2,3]. In the Iberian Pyrenees, anthropization initiated during the Bronze Age (∼3.7 cal kyr BP) in the southern lowlands [4] and progressed northward following an altitudinal trend characterized by rates of 40 m elevation per century, on average [5]. Therefore, climate was a major environmental forcing for Pyrenean vegetation during the LGM-Holocene transition.

Globally, the Lateglacial has been defined in Greenland ice cores by a stadial-interstadial succession that characterized the transition from the LGM to the Holocene, which is known as the INTIMATE (INTegration of Ice-core, MArine and TErrestrial record) event stratigraphy [6] (Fig. 1). The classical sequence, which serves as a worldwide stratigraphic reference, includes the stadial GS-2.1a or Oldest Dryas (OD) (17,500-14,700 cal yr BP), the interstadial GI-1 or Bølling/Allerød (B/A) (14,700-12,900 cal yr BP) – which has been subdivided into minor events GI-1a to GI-1e – and the stadial GS-1 (12,900-11,700 cal yr BP) or Younger Dryas (YD), which was the last Pleistocene cooling reversal that preceded the Early Holocene (EH) warming.

**Figure 1.**
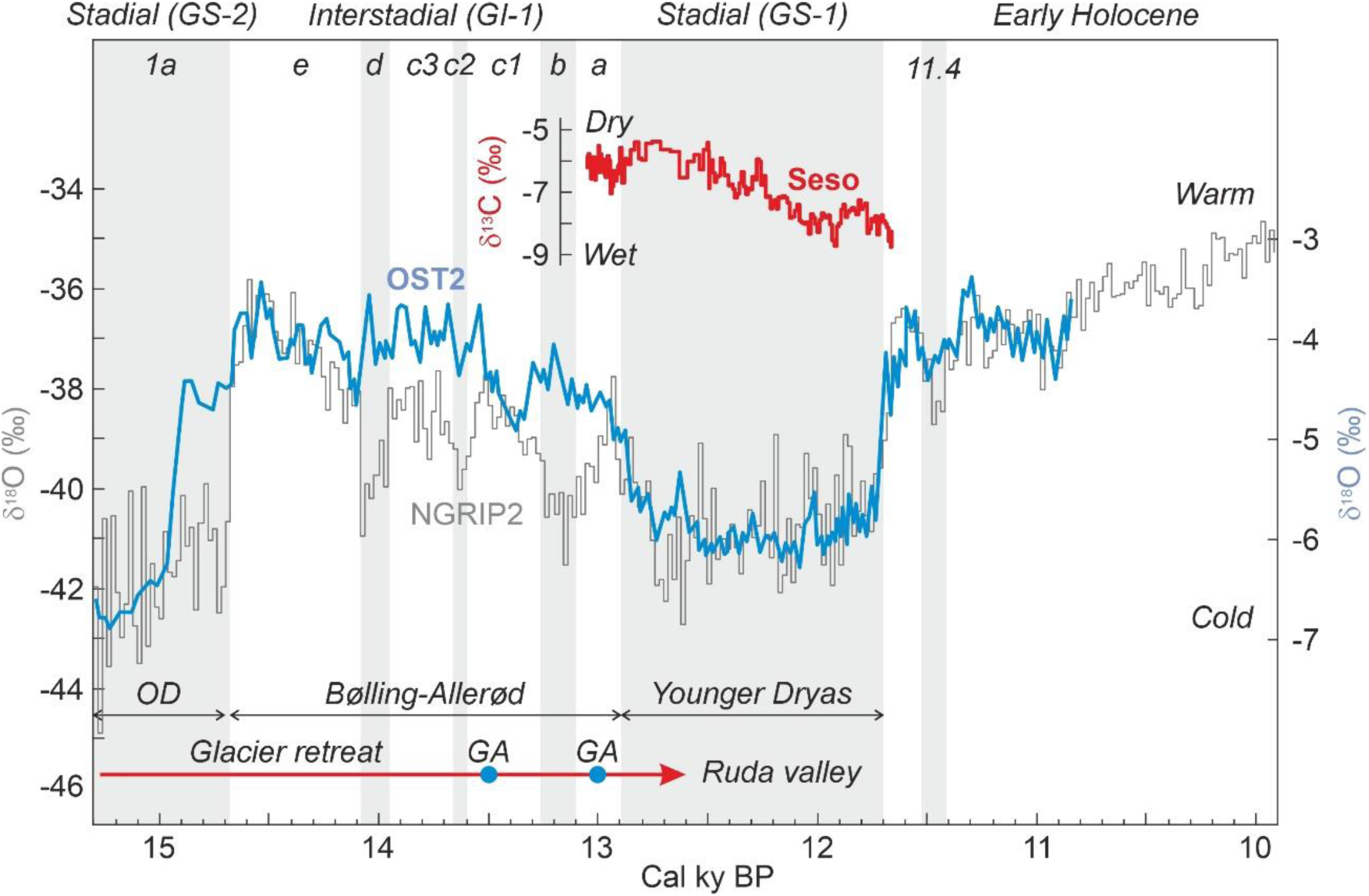
Lateglacial-Early Holocene stratigraphy based on Greenland ice core NGRIP2 [6] compared with the speleothem records from Ostolo cave (OST2), on the westernmost Pyrenean end [7] and Seso cave, on the central Pyrenees [8] (Fig. 2). The glacier dynamics, as reconstructed in the Ruda valley (parallel to the Aiguamòg valley, where the record analyzed in this study lies), is indicated for reference [9]. GA, glacier advance; OD, Oldest Dryas.

In the Pyrenees and surrounding lowlands, the Lateglacial-Early Holocene (LGEH) palynostratigraphic and paleovegetational trends are relatively well known on the northern side but largely unknown on the southern flank (Fig. 2). A first synthesis from the northern French slope [10] reported that the Lateglacial began with the expansion of *Betula*, the decrease in Chenopodiaceae and the decrease/disappearance of *Ephedra* at the initial Bølling. In some localities, *Juniperus* and *Pinus* also increased. Open cold/dry steppe environments were represented by *Artemisia*, which remained relatively constant, and increasing amounts of Poaceae, Apiaceae and *Rumex*. In drier Mediterranean areas (eastern sector), steppe assemblages were dominated by *Artemisia*, whereas in wetter Atlantic areas (western sector), the dominant were the Poaceae. A manifest *Pinus* increase coeval with a *Juniperus* decrease was recorded at the B/A transition, when *Quercus* began to establish and *Corylus* was still absent. During the YD, steppe elements (*Juniperus*, *Artemisia*, Chenopodiaceae, *Ephedra*) expanded again, while forest trees (*Betula*, *Pinus*) declined. This YD vegetation opening was more pronounced in the Mediterranean area due to the occurrence of drier conditions. At the end of the YD, *Artemisia* declined and *Ephedra* disappeared due to the onset of Holocene warming, characterized by the massive colonization of the Pyrenees by deciduous forests (*Betula*, *Corylus*, *Quercus*).

**Figure 2.**
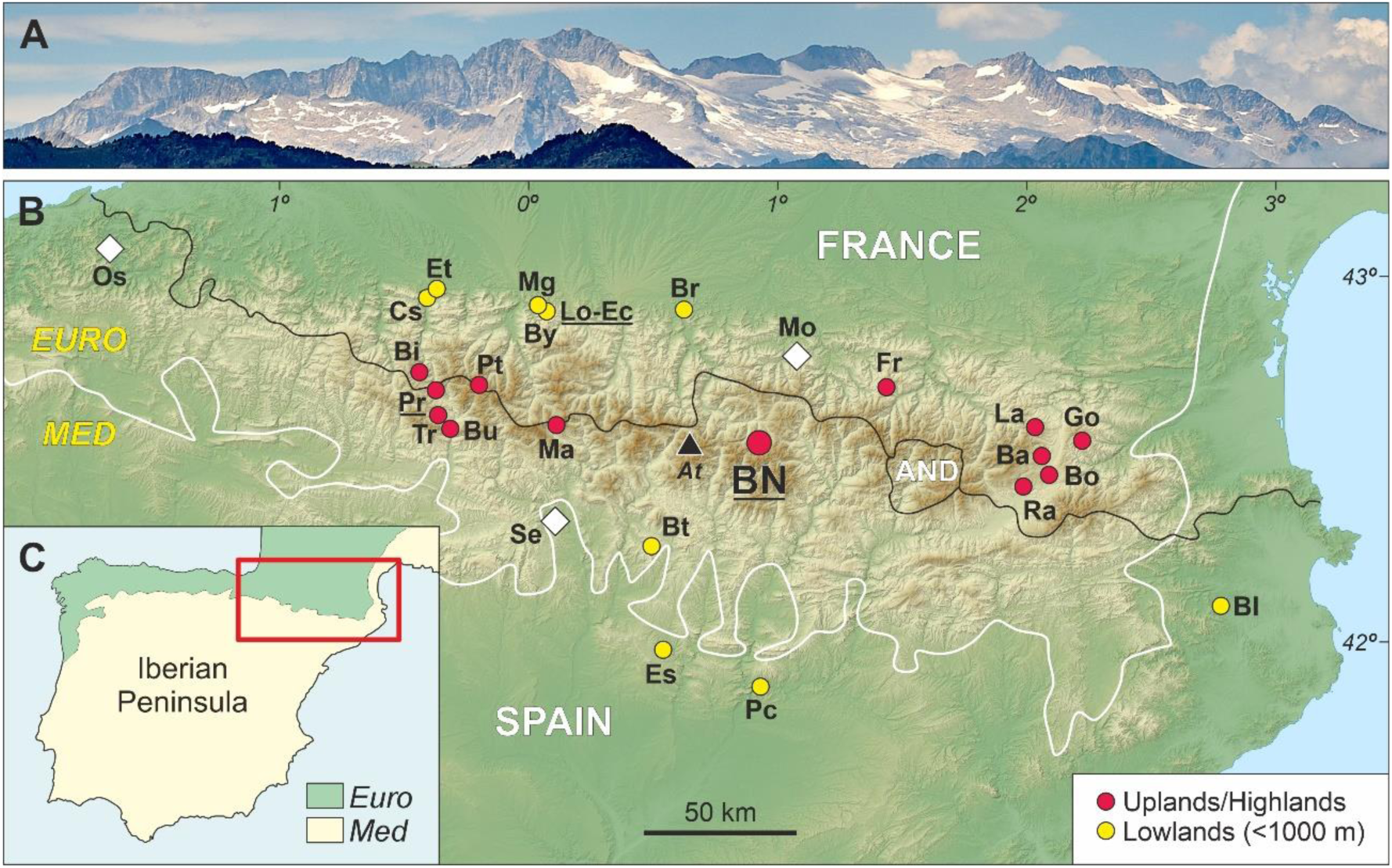
The Pyrenean range within the Iberian Peninsula. A) The Iberian Maladeta Massif, with the highest Pyrenean peak (Aneto; 3404 m) in the center, as seen from near the Bassa Nera (1890 m) coring site (Photo: V. Rull). B) Topographic map of the Pyrenees indicating the Bassa Nera coring site (BN) and other localities of the Pyrenees and the surrounding lowlands with Lateglacial pollen records. The locations of the Ostolo (Os), Seso (Se) and Moulis (Mo) paleoclimatic records are indicated by white diamonds, and the highest peak of the Pyrenees (At, Aneto) is marked by a black triangle. The black line is the boundary between Spain and France, which represents the approximate boundary between the northern and southern Pyrenean sides (AND, Andorra), and the white line is the boundary between the Eurosiberian and Mediterranean biogeographical regions. Lateglacial sites: Ba, Balcère; Bi, Bious; Bl, Banyoles; Bo, Borde; Br, Barbazan; Bt, Bentué; Bu, Búbal; By, Biscaye; Cs, Castet; Ec, Ech; Es, Estanya; Et, Estarrès; Fr, Freyxinède; Go, Gourg-Nègre; La, Laurenti; Lo, Lourdes; Ma, Marboré; Mg, Monge; Pc, Parco; Pr, Portalet; Pt, La Pouretère; Ra, Racou Tr, Tramacastilla (details and references in Appendix 1). Sites with charcoal records as fire proxies are underlined. C) Map of the Iberian Peninsula showing the location of the Pyrenees (red box). The Mediterranean (Med) and Eurosiberian (Euro) biogeographical regions are indicated by pale yellow and green areas, respectively.

Further high-resolution studies on some localities of the easternmost French Pyrenean side recorded an expansion of steppe elements (*Artemisia*, *Ephedra*, Poaceae, Apiaceae, Caryophyllaceae and others) and a decline in *Pinus* during the OD stadial [11]. The same studies documented a regional expansion of *Juniperus*, *Betula* and *Salix* during the initial B/A and a significant *Pinus* increase at the end of this interglacial. The YD stadial was characterized by a sudden decrease in *Pinus* and *Betula* and the expansion of steppe elements, especially *Artemisia*. The incoming warmer conditions during the Early Holocene were marked by a significant increase in *Betula* followed by *Quercus* and *Corylus*, which marked the initial colonization of the region by temperate forests. According to Reille & Lowe [11], oscillations in *Pinus* during Lateglacial times would have been largely due to fluctuations in long-distance pollen transport from low elevations; however, the local presence of *Pinus* forests during the Early Holocene was supported by the presence of stomata of *Pinus uncinata*. In general, increases in pollen from forest trees are considered to be indicative of forest expansions from neighboring refugia, which would have survived the last glaciation on sites with particular microclimatic conditions. The studies refined the existing chronologies and allowed a more precise dating of significant events such as the maximum of *Artemisia* (∼18.1 cal kyr BP), the rise of *Betula* (∼14.7 cal kyr BP), *Quercus* (∼12.3 cal kyr BP) and *Corylus* (∼11.5 cal kyr BP), or the first decline of *Betula* (∼12.8 cal kyr BP), among others [12].

A more recent study identified similar trends [13], but the low resolution of the LGEH sequence and the existence of a single radiocarbon date in this interval prevent detailed correlations. A more recent multicentennial record identified the following succession: *Juniperus*-*Artemisia* (OD) – *Pinus*-*Betula*-Poaceae (B/A and YD) – *Pinus*-*Betula*-*Quercus*-*Corylus* (Early Holocene) [14]. This work included charcoal analysis and documented no fire activity during the OD, low to medium fire incidence during the B/A and the YD, and a sudden fire increase during the Early Holocene. These patterns were explained in terms of climate-vegetation interactions, with the former as the main driver and the latter as a modulator of the amplitude of fire regimes. In this way, the abrupt fire increase at the beginning of the Holocene was linked to the expansion of temperate deciduous woodlands.

On the southern Pyrenean side, only four LGEH palynological sequences are available, all of them on the western side (Fig. 2). Three of these records (Marboré, Búbal and Tramacastilla) are of multicentennial resolution, and the LGEH stadials/interstadials are not well defined palynostratigraphically. In addition, in the Búbal and Tramacastilla records, chronological correlations among similar palynological zones are problematic despite their proximity (barely 6 km distant), which was attributed to methodological dating issues [15]. The fourth record, Portalet, is well bracketed chronologically and of higher resolution but incomplete, as the YD is missing by a sedimentary gap, which was attributed to the permanent freezing of the Lateglacial paleolake [16]. The lack of highland YD palynological records is the norm on the southern side, where this climatic reversal is represented only in the lowlands by cold-steppe elements (notably *Artemisia*, Poaceae, Chenopodiaceae and *Juniperus*) [review in Ref. 17]. Therefore, a chronologically well-bracketed complete and continuous LGEH palynological sequence of centennial or higher resolution for the Iberian Pyrenees is lacking.

In general, increases in pollen from forest trees (*Pinus*, *Betula*, *Corylus*, *Quercus*) similar to those documented for the northern Pyrenean slopes occurred in the Iberian Pyrenees during warming phases (B/A, EH). These increases are believed to reflect temperate forest expansions from nearby glacial refugia situated in the southern lowlands (<800 m) as part of a highly diverse mosaic landscape composed of conifer forests, mixed pine-oak forests, parklands, grasslands and xerophytic steppes [18,19]. Similar to the northern side, LGEH Pyrenean records from the Iberian side usually lack charcoal records as fire proxies, except for the Portalet sequence, where Lateglacial wildfire incidence was lower than in the Early-Mid Holocene, probably owing to the combination of higher temperatures and increased biomass fuel due to the expansion of deciduous forests since 10 cal kyr BP [20]. Other palynological parameters and proxies unusual on both northern and southern Pyrenean slopes are pollen accumulation rates (PAR), as a proxy for plant cover [21], and non-pollen palynomorphs (NPP), which have demonstrated to be very useful for environmental reconstruction [22,23].

This paper presents the Bassa Nera record, which is the first well-dated complete and continuous (gap-free) Lateglacial-Early Holocene (∼15,200-10,000 cal yr BP) palynological record of vegetation change at centennial resolution from the central Iberian Pyrenees, including PAR, charcoal and NPP analyses. The main Holocene trends (10 cal kyr BP onward) in the development of the Bassa Nera vegetation have already been studied [24–27], but the LGEH ecological history remains unknown. The main aims of this study are (i) to provide a standard LGEH palynological sequence for the central Pyrenees, which may serve correlation and dating purposes; (ii) to characterize the response of vegetation to local and regional climatic shifts in the absence of human disturbance, which may be useful to disentangle natural and anthropogenic drivers of ecological change; (iii) to document the fire regime history and its potential climatic and vegetational drivers; and (iv) to situate the results obtained in a regional context by comparing them with former studies on the northern and southern Pyrenean sides, including the surrounding lowlands. In addition to providing clues on the origin of ecosystems and their sensitivity to natural forcings, knowing the responses of preanthropic vegetation to climate changes is useful to anticipate the potential responses of natural communities to future climatic change.

## 2. Study area

### 2.1. Present conditions

Bassa Nera is a small pond (ca. 100 m long, 60 m wide and 5 m deep) located in a small depression known as Pletiu dera Muntanheta, situated in the headwaters of the Aiguamòg valley (upper Garona basin), at 42° 38’ 17” N - 0° 55’ 27” E and 1890 m elevation. Although it belongs to Spain (Val d’Aran), the site is located on the northern Pyrenean slope, near the Pyrenean ridge. In some maps, the pond is also called Estanh dera Muntanheta, but we prefer to use Bassa Nera because this is the most utilized name in recent botanical and paleoecological publications. The pond is in its later infilling stages and is surrounded by a mire. The site is situated in the peripheral protection zone of the National Park “Aigüestortes i Estany de Sant Maurici” (PNAESM). The Pletiu dera Muntanheta is of glacial origin and is excavated on the granodioritic bedrock that is part of the Carboniferous-Permian Maladeta batholith [28,29]. The climate is alpine with oceanic influence, and the regional 0 °C isotherm is estimated to be at 2950 m elevation [30]. The nearest meteorological station is Port de la Bonaigua, situated approximately 6 km NE and at 2072 m elevation, 182 m above Bassa Nera. The annual average temperature for the 1990-2020 period is 3.34 °C, and the total annual precipitation is 1218 mm. Minimum temperatures (-2.9 to -3.2 °C) are recorded in January and February, and maxima (11.6 °C) correspond to July and August. Total precipitation shows a bimodal pattern with maxima (131-137 mm) in May and November and minima (70-79 mm) in February and July.

Biogeographically, the Bassa Nera is located in the southern margin of the Eurosiberian region, characterized by wet temperate Atlantic macrobioclimates and dominated by plant species distributed across central and northern Europe. This contrasts with the Mediterranean biogeographic region, under the influence of Mediterranean climates with at least two consecutive arid months during the warmest season (summer). The Mediterranean biome dominates most of the Iberian Peninsula (Fig. 2) and is characterized by plant species distributed across southern Europe and northern Africa [31].

A study of plant communities along the Aiguamòg valley ranging between approximately 1300 m and 2600 m elevation, which included the Pletiu dera Muntanheta, recognized the following vegetation stages: mixed forests of *Pinus sylvestris* and *Betula* (1300-1600 m); conifer forests of *Pinus mugo* and *Abies* (1600-2000 m); mosaic vegetation of *Pinus mugo*, *Rhododendron* and *Nardus* (2000-2250 m); and alpine meadows/bogs with *Festuca*, *Nardus*, *Carex*, *Luzula* and *Juncus* [32] (Fig. 3). In this transect, the Bassa Nera lies within the *Pinus*-*Abies* forest belt, ∼350 m below the treeline, which is situated at 2250 m. A further study specifically of the Pletiu dera Muntanheta identified seven plant associations [33]. The pond margins are characterized by *Sphagnum* peats, *Carex*-*Potentilla* associations, or *Sphagnum*-*Carex* carpets, depending on the local features. The surrounding areas are covered by marshes and grasslands dominated by *Carex*, *Scirpus* or *Succisa*, with scattered *Sphagnum* hummocks. *Pinus*-*Abies* forests with a *Rhododendron*-dominated understory grow on encircling slopes. The same study highlighted the uniqueness of the site in terms of plant richness in comparison with other Pyrenean mires and attributed this fact to the combination of hydrological and water chemistry variability.

**Figure 3.**
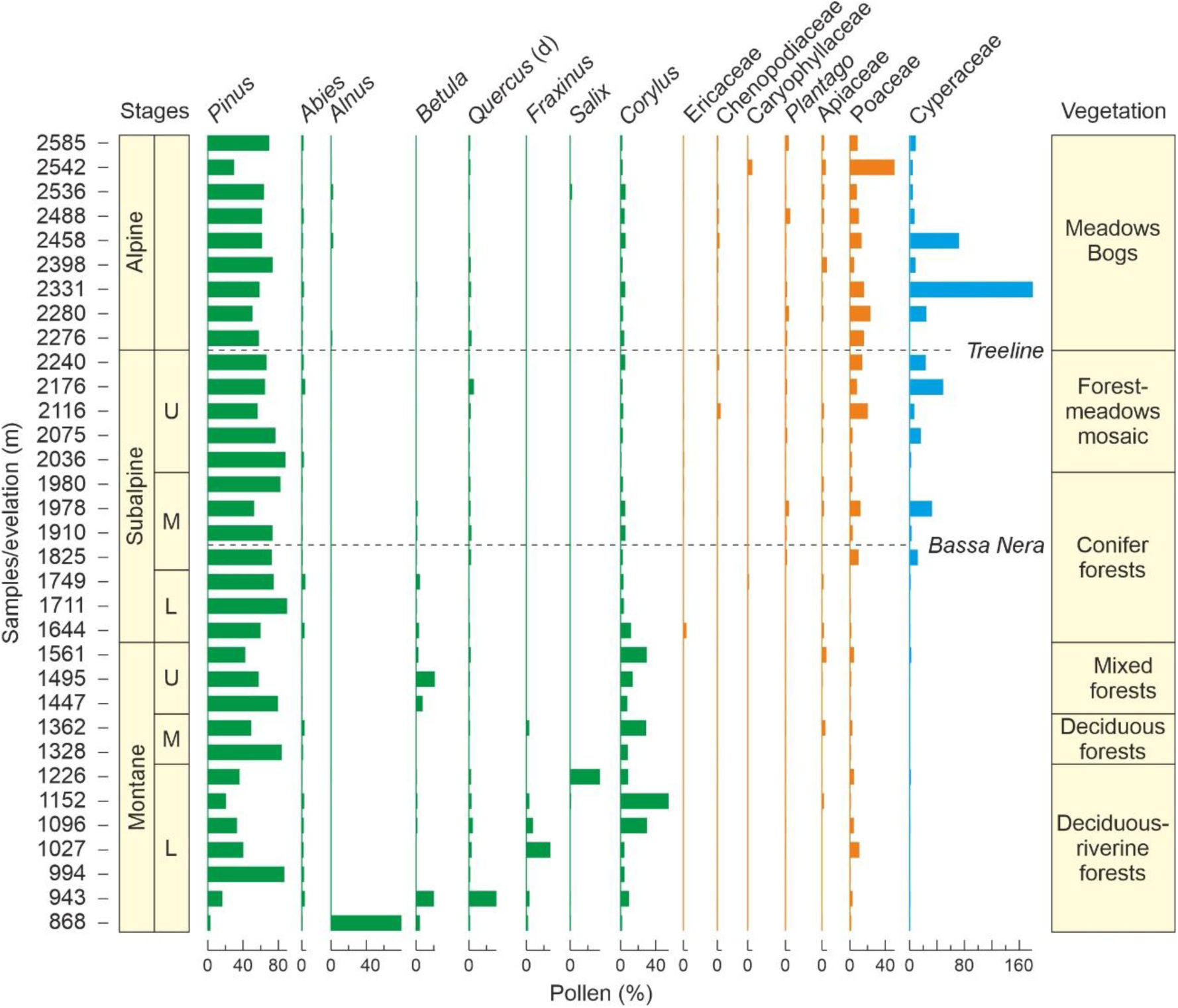
Modern pollen spectra along an elevational transect including the Bassa Nera coring site [32]. Only major components (>2% of the total) are shown. Forest trees are in green, and shrubs/herbs are in orange. Cyperaceae (blue) were not included in the pollen sum.

### 2.2. Previous studies

The Pletiu dera Muntanheta was included in a modern-analog study developed on an altitudinal transect (ca. 870-2600 m elevation) along the Aiguamòg valley. This survey was aimed at establishing qualitative and quantitative relationships between extant vegetation patterns and modern pollen sedimentation, which may be useful for the interpretation of past pollen records in terms of vegetation change [32]. The lower montane vegetation belt (800-1300 m) was characterized by *Alnus*, *Betula*, deciduous *Quercus*, *Fraxinus* and *Salix* pollen, which is absent in the upper levels (Fig. 3). In the middle and upper montane levels (1300-1600 m), the pollen assemblages were almost exclusively composed of *Pinus*, with lower amounts of *Corylus*. The subalpine belt (1600-2300 m; below the treeline) was absolutely dominated by *Pinus*, with low percentages of grass and sedge pollen, whereas the pollen of deciduous trees from montane belts was virtually absent. The situation was similar in the alpine belt (2300-2600 m; above the treeline) but with higher amounts of herbaceous plants, which may be locally more abundant than *Pinus*. The best altitudinal indicators were found in *Fraxinus* and *Salix*, whose pollen is restricted to the montane and subalpine belts, respectively. *Betula* and *Corylus* pollen was restricted to the montane belt, together with deciduous *Quercus*, *Tilia*, *Alnus*, *Buxus* and Brassicaceae. Pollen types almost restricted to the alpine belt were *Plantago*, *Calluna*, Poaceae, Asteraceae and Cyperaceae, which may serve as treeline indicators. Interestingly, the pollen of *Olea* and evergreen *Quercus* have their maxima in the subalpine belt, although their parent plants are characteristic of lowlands and were not found in the vegetation gradient analyzed, suggesting efficient upward wind dispersal. In the same transect, the modern sedimentation of NPP was studied to unravel their potential as paleoecological indicators [26]. These authors found that NPP assemblages are influenced not only by elevation but also by local moisture conditions, landscape openness and human pressure, especially grazing.

Previous studies in the Pletiu dera Muntanheta based on cores retrieved in the mire surrounding Bassa Nera recorded the vegetation shifts of the last ∼10,000 years. These studies demonstrated that human pressure was low until the Late Bronze Age. Indeed, the first evidence of grazing dates to 7300 cal yr BP, and the oldest cereal records were found at ∼5190 cal yr BP, but general landscape anthropization did not occur until 3150-2650 cal yr BP [25]. No archaeological evidence of human occupation of the PNAESM prior to the Holocene is available, and Neolithization began by ∼7250 cal yr BP at lower elevations (<1600 m) and ∼5350 cal yr BP at higher elevations [34].

No paleoecological studies are available for the LGEH, but a recent glaciological survey near Bassa Nera provides the paleoclimatic context for the interpretation of the palynological studies presented in this paper. LGEH chronology and paleoclimatology were resolved in detail in the adjacent Ruda valley using ^10^Be cosmic-ray exposure dating of moraine boulders and polished surfaces, combined with glacier equilibrium-line altitude calculations [9]. After the LGM, a general glacier retreat occurred that exposed areas at elevations of approximately 1860-1900 m between 15 and 14 cal kyr BP. The Bassa Nera lies within this elevational range; thus, it can be inferred that the Pletiu dera Muntanheta was covered by glaciers until ∼15 cal kyr BP, when our pollen record begins (Fig. 1). Therefore, this record may be the oldest possible to be retrieved in the Bassa Nera sediments and represent periglacial environments. The deglaciation trend continued until the YD and the Holocene but was interrupted by two glacier advances/stillstands at 13.5 and 13.0 cal kyr BP, represented by moraine systems situated at 2080 m and 2190 m elevation, which means that the glacier fronts were 190 m and 300 m above Bassa Nera, respectively. The deglacial trend exposed the surface of the highest cirque floors of the Ruda valley (2300-2500 m elevation) by 12.7-12.6 cal kyr BP, at the onset of the YD [9].

## 3. Methods

### 3.1. Coring and dating

The Bassa Nera core used in this study (PATAM12-A14) was retrieved in July 2014 with a 50-mm diameter Livingstone corer. This core consisted of 10 drives (D1 to D10) for a total length of 845 cm (1102 cm once corrected for compression). This study is focused on the basal sequence (drives D7 to D9), which contains the LGEH interval. Eight samples for radiocarbon dating were taken in this interval and submitted to acid/base digestion (KOH, HCl, HF) to remove the rejuvenating effect of eventual incorporation of younger plant material and/or humic acids percolating through the peat profile [35]. AMS radiocarbon dating was performed at the Radiochronology Lab of the University Laval (Canada). Calibration was performed with *CALIB 8.2* and *IntCal 20* [36]. The Bayesian age-depth model was built with the software BACON [37].

### 3.2. Palynological analysis

For pollen analysis, 51 samples (2 cm^3^) were taken with a volumetric syringe at regular intervals of approximately 100 years once the age-depth model was known (Fig. 1). These samples were digested with NaOH for disaggregation, HCl for carbonate removal and HF for silicate removal and submitted to centrifugation with Thoulet solution (density 2 g/cm^3^) for mineral separation [38]. Two tables of *Lycopodium* spores (Batch 280521 291; 13,761±448 spores/tablet) were added to each sample before processing [39]. Processing was performed at the Archaeobotany Lab of the Catalan Institute of Human Paleoecology and Social Evolution (IPHES). Pollen/spore identification was based on usual pollen catalogues for the region [40,41] and the author’s (VR) own reference collection, which contains over 700 species/subspecies from the Iberian flora. NPP identification was based on our previous research [26,27] and the NPP Image Database (https://non-pollen-palynomorphs.uni-goettingen.de/; last visited 3 May 2023). Most fungal spores were not identified, but special attention was placed on those considered to be indicators of herbivory and therefore of human impact, such as *Sporormiella*, *Sordiaria* and *Podospora* [42–45].

Counting was conducted until a minimum pollen sum of 300 grains, stabilization of confidence intervals and saturation of diversity [46]. In a number of samples, *Pinus* was superabundant, and its counts were stopped at 100-150 grains and further extrapolated using *Lycopodium* spores. In those cases, the above statistical criteria were applied to the remaining pollen types. The pollen sum included all pollen types with the exception of aquatic and semiaquatic plants (Cyperaceae, *Typha*, *Ludwigia*). Paleoecological interpretation was based on the available modern-analog studies for both pollen/spores and NPP developed on the elevational transect that includes the study site [26,32]. Pollen accumulation rates (PAR) are used as a proxy for vegetation cover [21]. It is important to stress that pollen is not used as a climatic proxy, as usual in most Pyrenean literature. This is due to the lack of sufficient taxonomic resolution (usually at the genus or family level) and to the fact that such a procedure may lead to circularity when analyzing the response of vegetation to changing climates. Paleoclimatic trends were inferred from pollen-independent proxies such as the OST2 (temperature) and Seso (moisture) speleothem records (Figs. 1 and 2). Unfortunately, independent paleoclimatic proxies for hydrological balance are available only for the Younger Dryas [8,47]. Charcoal particles above 5 μm were also recorded and used as fire proxies. Two size classes of these particles were considered, above and below 150 μm, as proxies for fires around the coring site and regional fires, respectively [48].

Zonation of pollen diagrams was performed using the optimal splitting by information content (OSIC) and broken-stick methods [49]. Cluster analysis used the Pearson linear correlation coefficient (r) and the unweighted pair-group average (UPGMA) clustering algorithm [50]. Plotting and zonation were performed with *Psimpoll 4.27* [51], and statistical analyses were carried out with *MVSP 3.22* [52] and *Past 4.12* [53].

## 4. Results

### 4.1. Lithology and Chronology

The full core was subdivided into five sedimentary units, of which three (U3 to U5) were present in the LGEH interval studied here. These units are characterized by intercalated clay, silt and sand layers of varied appearance and composition. U3 was composed of gray clays and fine sands with plant macroremains, whereas in U4, the alternating laminae consisted of brown clays and ochre silts/sands with low organic matter content, and U5 was dominated by brown greenish clay layers with no apparent organic matter and occasional gravel and silt/sand beds (Fig. 4). Radiocarbon dating (Table 1) showed that the interval studied contained a complete and continuous (i.e., free from sedimentary gaps) LGEH sequence that lasted approximately 5000 years, from 15.3 to 10.4 cal kyr BP, and included the GS-2 (OD) stadial, the GI-1 (B/A) interstadial, the GS-1 (YD) stadial and the EH warming (Fig. 1). The average sediment accumulation rate (r) was 0.63 mm yr^-1^, but it was far from homogeneous, showing minimum to intermediate values (0.29-0.82 mm yr^-1^) in U3 and U5 and maximum values in U4, especially in its lower part (0.90-1.60 mm y^-1^). This acceleration event occurred in the first part of the YD (12.8-12.3 cal kyr BP) and is abbreviated here as YDsa.

**Figure 4.**
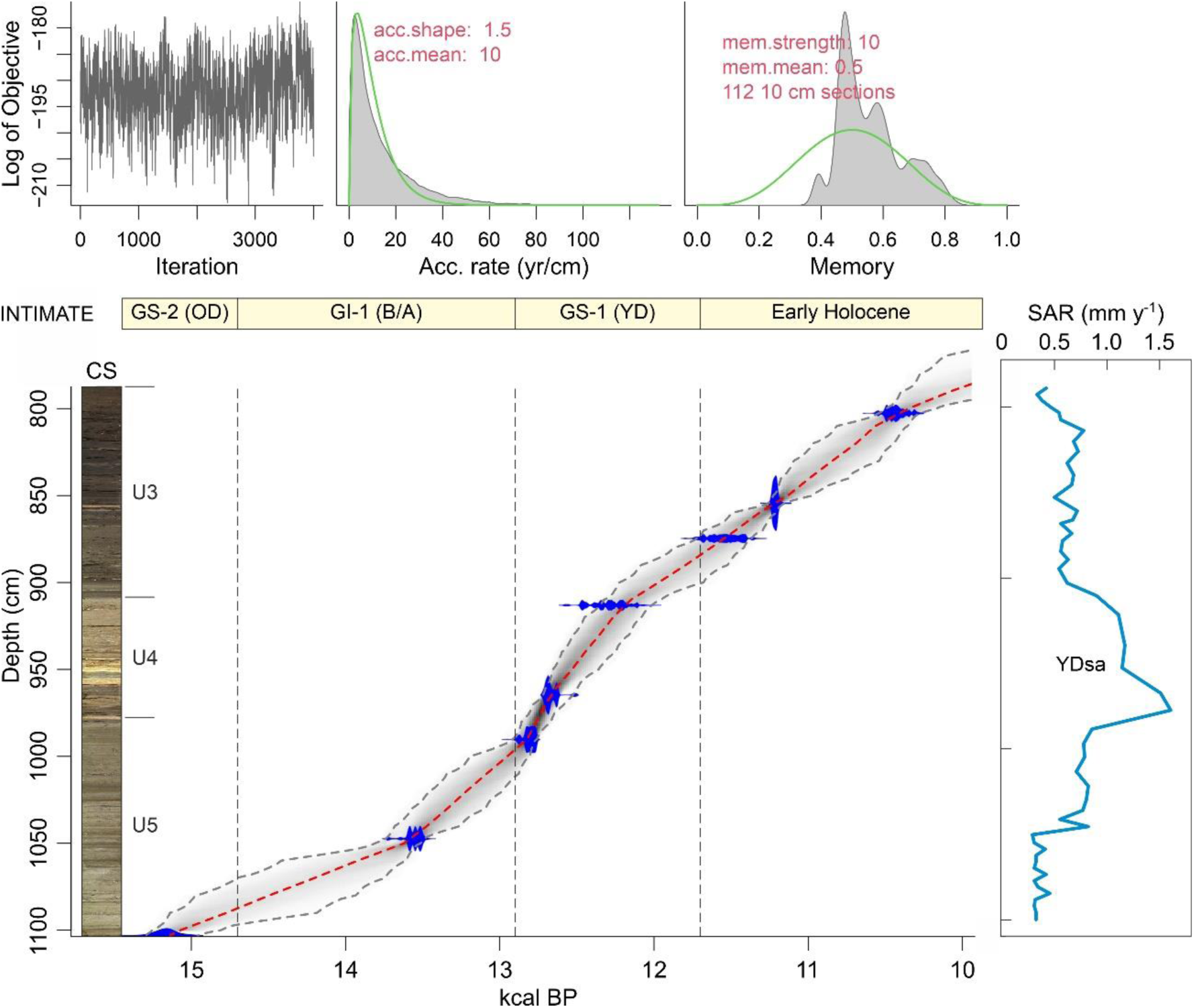
Lithology, BACON age-depth model [37] and sediment accumulation rates (SAR) of the core analyzed in this work. The INTIMATE stadial/interstadial stratigraphy (Fig. 1) is also represented for reference. CS, core scan; U3-U5, sedimentary units; YDsa, Younger Dryas sedimentation acceleration.

**Table 1.** Radiocarbon dates of the basal part of core PATAM12-A14 using pollen residues [35]. <u>Dating</u>: Radiochronology Lab, University Laval, Canada (ULA). <u>Calibration</u>: *CALIB 8.2* and *IntCal 20* [36]. Sedimentation rates (r) refer to the intervals after the corresponding age values; for example, 0.35 mm yr^-1^ corresponds to the interval 15,167-13,549 cal yr BP.

| Lab code | Drive | Sample | Depth (cm) | <sup>14</sup>C yr BP | Cal yr BP (2σ) |
| --- | --- | --- | --- | --- | --- |
| ULA-10114 | D7 | 63-64 | 802 | 9260±20 | 10,300-10,557 |
| ULA-10115 | D7 | 104-105 | 854 | 9785±20 | 11,191-11,241 |
| ULA-10116 | D8 | 19-20 | 874 | 10,040±20 | 11,358-11,727 |
| ULA-10117 | D8 | 51-52 | 913 | 10,410±20 | 12,102-12,482 |
| ULA-10118 | D8 | 94-95 | 964 | 10,635±20 | 12,621-12,716 |
| ULA-10119 | D9 | 32-33 | 990 | 10,915±20 | 12,757-12,882 |
| ULA-10120 | D9 | 76-77 | 1047 | 11,710±25 | 13,493-13,602 |
| ULA-10121 | D9 | 119-120 | 1103 | 12,715±25 | 15,049-15,281 |

### 4.2. Palynology

Globally, a total of 56 palynomorph types (33 pollen, 8 pteridophyte spores and 15 NPP) were identified. Total counts averaged 914 (804 pollen, 18 pteridophyte spores and 92 NPP), and the average pollen sum was 792. The pollen diagram was subdivided into seven pollen zones, which are described in chronological order. Pollen and spore percentages are shown in Fig. 5, NPP percentages are depicted in Fig. 6 and influx values are displayed in Fig. 7. Boundary ages, in 95% confidence intervals, were estimated using the age-depth model of Fig. 4. The most abundant NPP are depicted in Plates 1, 2 and 3.

**Figure 5.**
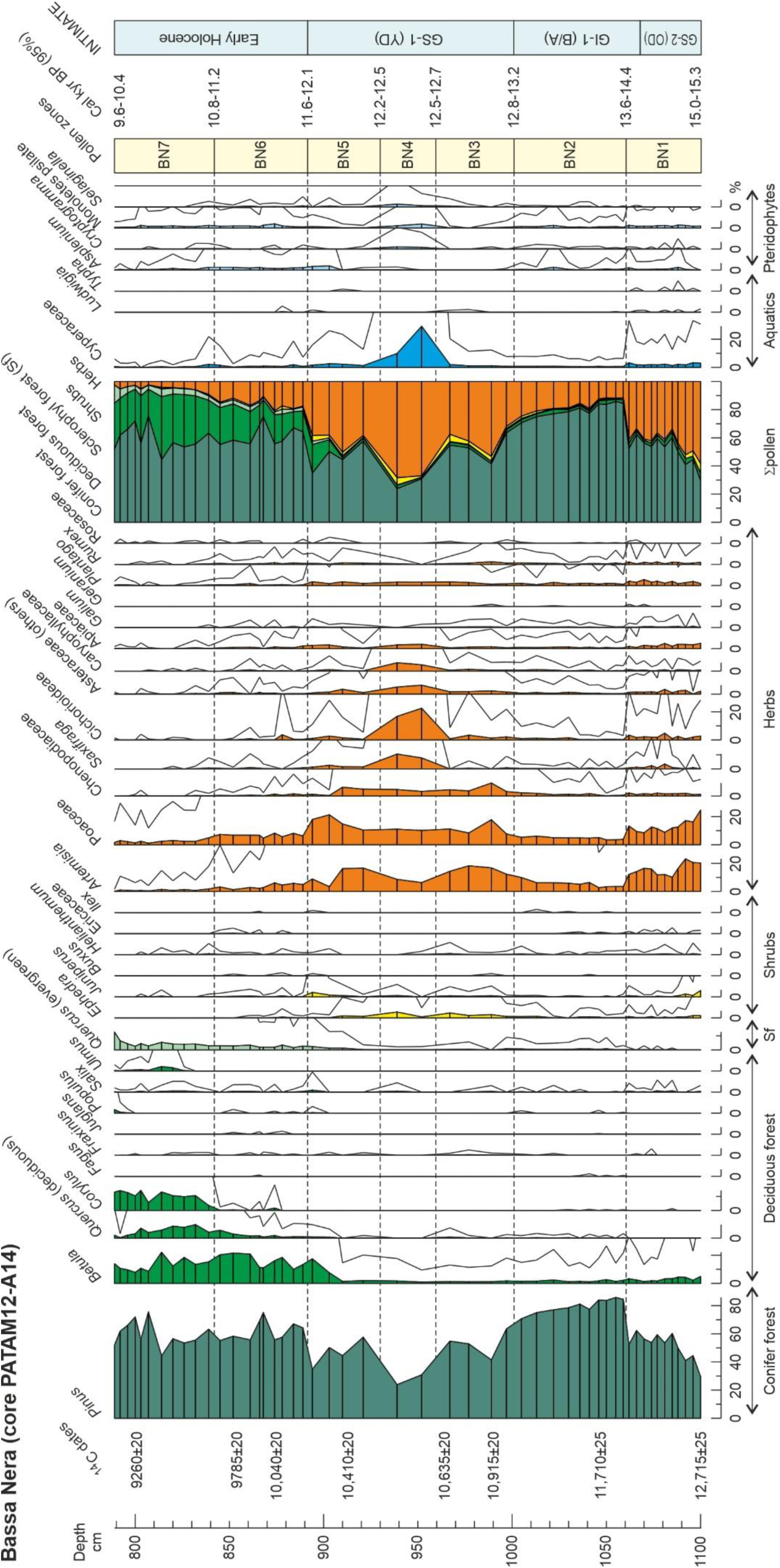
Percentage diagram of pollen and pteridophyte spores showing the pollen zones (BN1 to BN7) established using the OSIC and broken stick methods (Bennett, 1996). Solid lines are x10 exaggeration. Arboreal pollen is subdivided into conifer forest (Cf), deciduous forest (Df) and sclerophyll forest (Sf) elements; nonarboreal pollen is subdivided into shrubs (Sb) and herbs (Hs). Σp, pollen sum. Boundary ages of pollen zones are expressed as 95% confidence intervals obtained by BACON age-depth modelling (Fig. 4).

**Figure 6.**
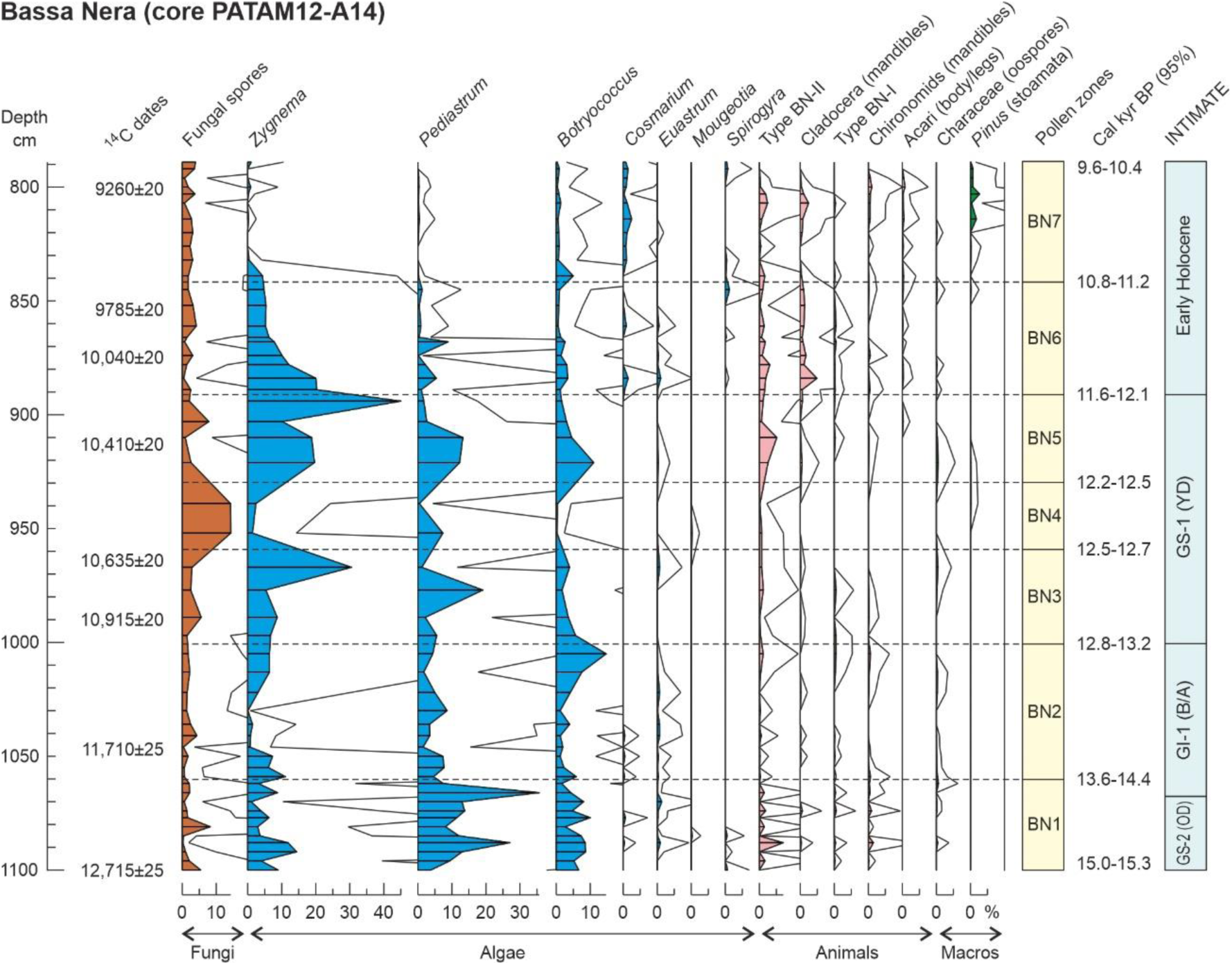
Percentage diagram of non-pollen palynomorphs (NPP) using the pollen zones defined in Fig. 5. Solid lines are x10 exaggeration. The most abundant NPP are depicted in plates 1, 2 and 3. Pollen zones and boundary ages as in Fig. 4.

**Figure 7.**
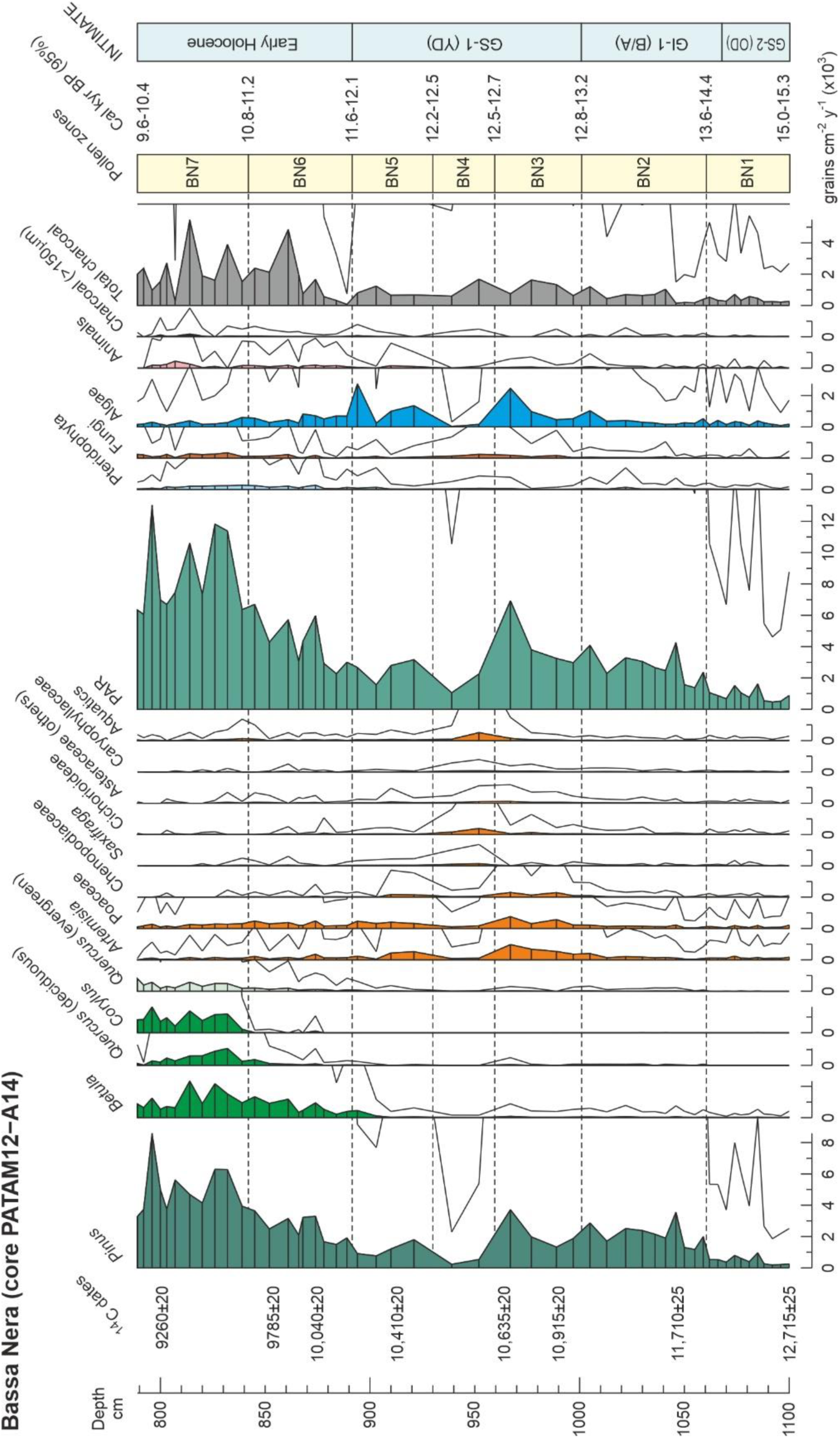
Summary influx diagram displaying the most abundant pollen types, the NPP groups and charcoal. Solid lines are x10 exaggeration. Pollen zones and boundary ages as in Fig. 4. PAR, pollen accumulation rates.

**Plate 1.**
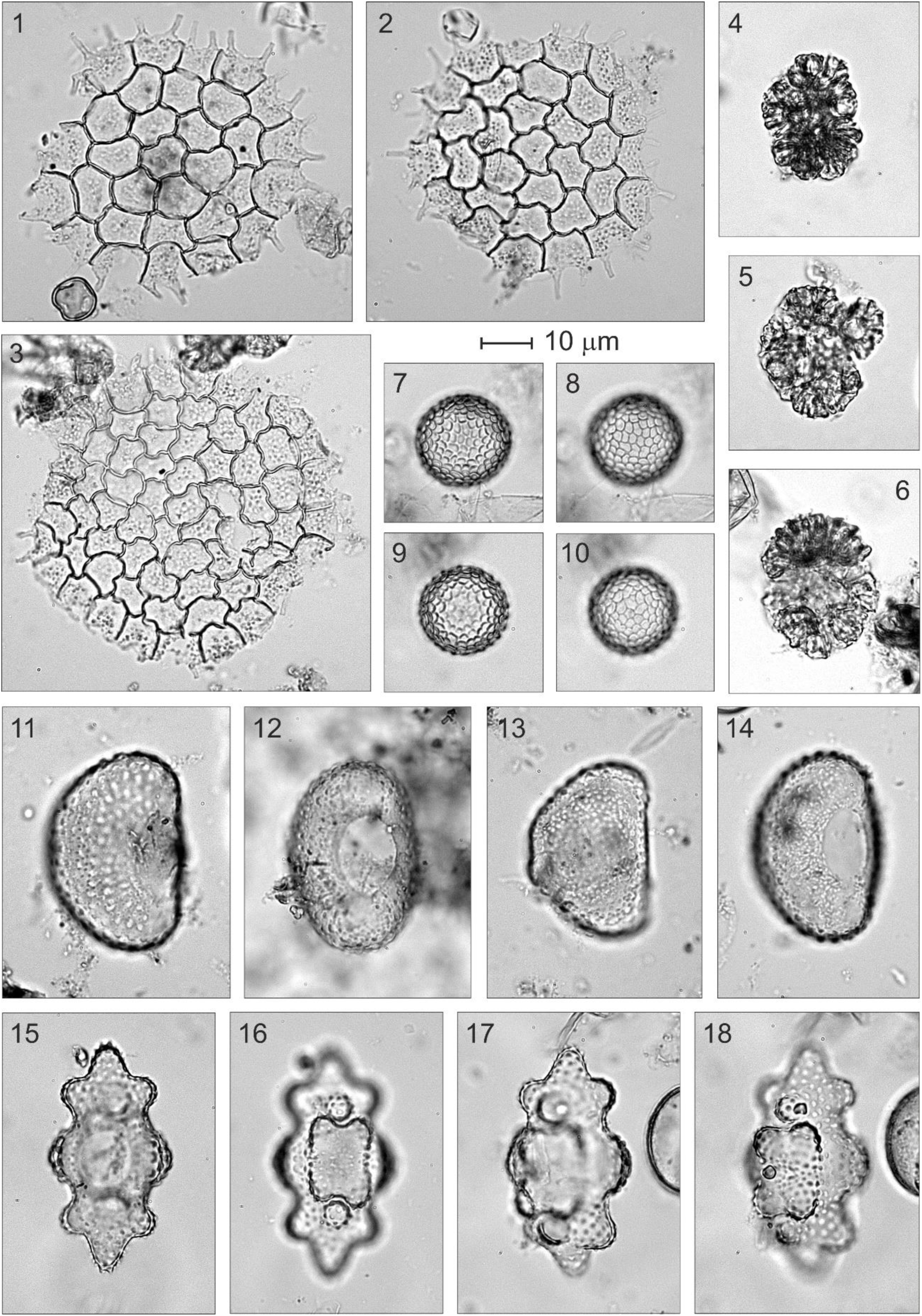
Algae remains. 1-3, *Pediastrum*; 4-6, *Botryococcus*; 7-10, *Zygnema*; 11-14, *Cosmarium*; 15-18, *Euastrum*.

**Plate 2.**
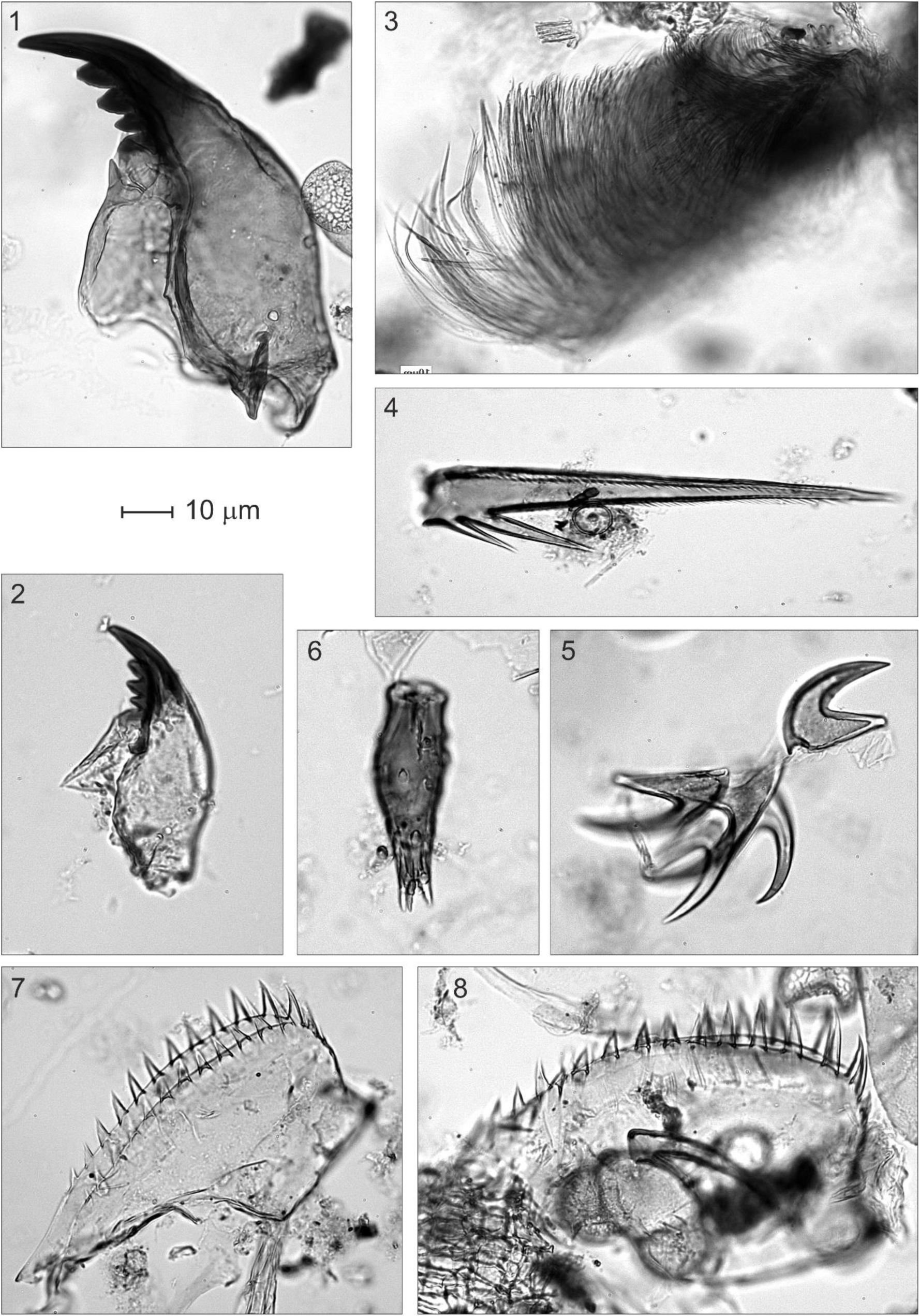
Animal remains. 1-2, Chironomids (mandibles); 3, Type BN-I; 4, Type BN-IIa; 5, Type BN-IIb; 6, Acari (legs); Cladocera (mandibles).

**Plate 3.**
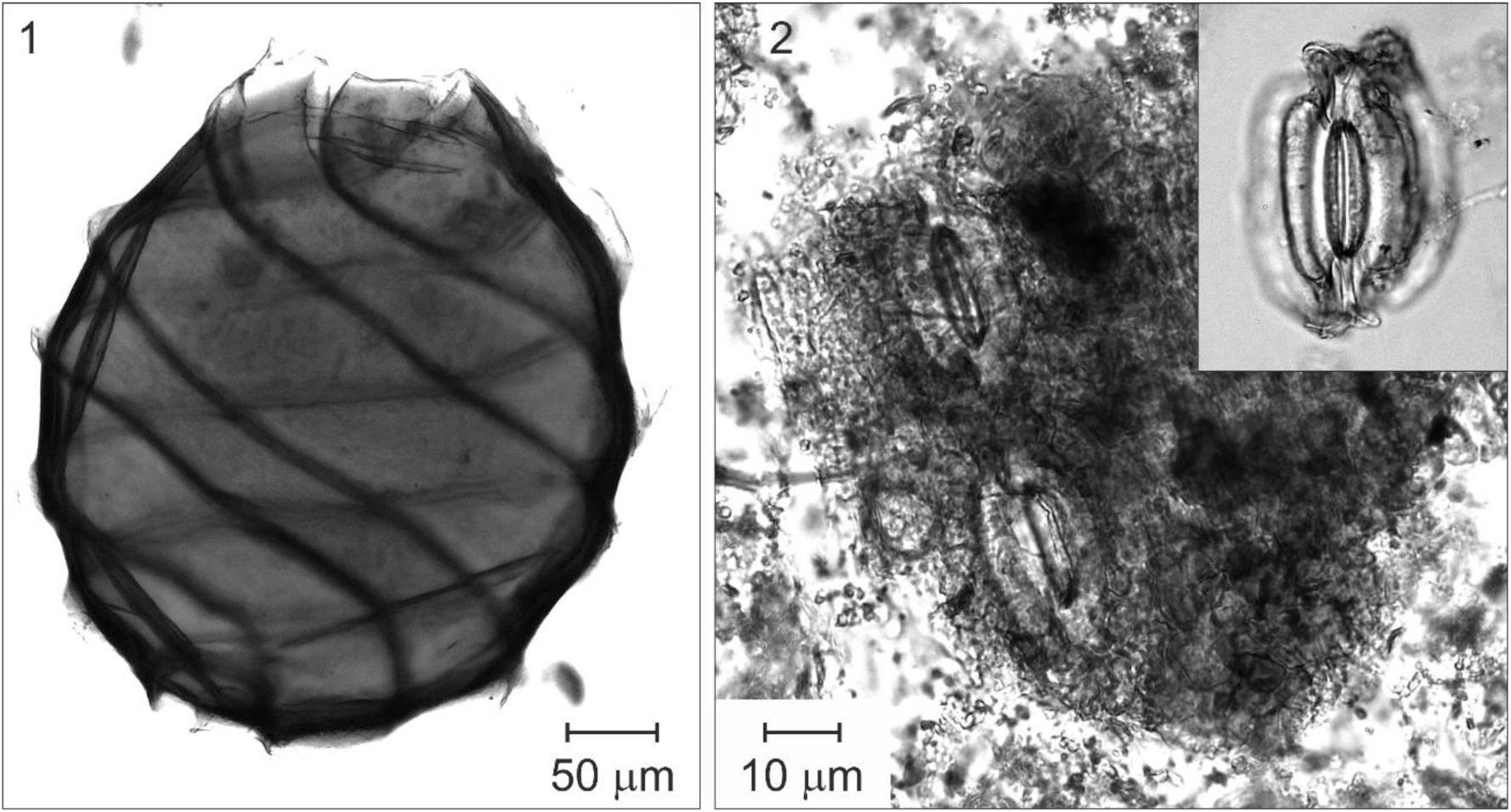
Plant macroremains. 1, Characeae (oospore); 2, Fragment of epidermis of a pine leaf with stomata and an isolated stoma (upper right corner).

#### Zone BN1 (1100 to 1061 cm; 14,961-15,300 to 13,635-14,445 cal yr BP; late GS-2/OD)

Codominance of arboreal and nonarboreal elements. The dominant tree was *Pinus* (>40%), followed by the herbs *Artemisia* and Poaceae (15-20% each), all other elements were absent or below 1% of the pollen sum. These dominance relationships were due to the general scarcity of pollen, as indicated by minimum PAR values, rather than to high abundances of major types. Low percentages of fungal spores (∼5%). NPP also scarce and dominated by *Pediastrum* (alga) and Type BN-II as the main animal remain. Minimum charcoal influx.

#### Zone BN2 (1061 to 1001 cm; 13,635-14,445 to 12,821-13,191 cal yr BP; late GS-2/OD to GI-1/BA)

Arboreal elements largely dominant (70-80%) due to the increase in *Pinus* (60-80%) and the decline in *Artemisia* and Poaceae (<10%). This dominance shift was due to an increase in *Pinus* influx, which was the responsible for a general increase of PAR values, whereas *Artemisia* and Poaceae remained at values similar to Zone BN1. General percentage decrease of NPP except fungi spores, which remained similar to the former zone. Algae (especially *Botryococcus*, which peaked at the end of the zone) and charcoal influx slightly increased upward.

#### Zone BN3 (1001 to 960 cm; 12,821-13,191 to 12,493-12,701 cal yr BP; early GS-1/YD)

Transitional zone representing a gradual reversal to values similar to BN1, except for PAR, which showed a maintained increase and peaked at the end of the zone. In this case, however, the increase in relative abundance of *Artemisia*, Poaceae and Chenopodiaceae was due to the actual increase in their pollen influx, reaching a maximum at the end of the zone. Among NPP, fungi displayed an increasing trend and algae significantly increased in both percentage and influx, especially *Zygnema* and *Pediastrum*, whereas *Botryococcus* decreased. Charcoal influx continued its slightly ascending trend.

#### Zone BN4 (960 to 930 cm; 12,493-12,701 to 12,164-12,524 cal yr BP; middle GS-1/YD)

Dominance of nonarboreal elements (∼70%) and significant decline in PAR values after the ending-BN3 peak. Apparent decrease in *Pinus*, and slight decline in *Artemisia*, Poaceae and Chenopodiaceae, in both percentage and influx. The situation was similar to BN3 but Cichorioideae, Cyperaceae and *Saxifraga*, experienced a slight influx increase, which resulted in a meaningful percentage increase, followed by other Asteraceae, Caryophyllaceae and Apiaceae. Fungal spores showed their maximum percentages (although this was not due to a conspicuous influx increase), whereas algae and animal remains remarkably declined reaching their minimum values, in both influx and percentage. Charcoal slightly decreased.

#### Zone BN5 (930 to 892 cm; 12,164-12,524 to 11,584-12,105 cal yr BP; late GS-1/YD)

Another transitional zone showing trends opposite to BN3, that is, increasing arboreal and decreasing nonarboreal pollen. PAR values did not recover completely and remained below BN3. First significant increase in *Betula* at the middle of the zone and further increase in both percentage and influx. Nonarboreal elements recovered the situation of BN3 due to the strong decline of Cichorioideae, Cyperaceae, *Saxifraga*, other Asteraceae, Caryophyllaceae and Apiaceae. Fungi declined to former BN3 percentage values and all algae, especially the dominant *Zygnema*, underwent significant increases, peaking at the end of the zone. Among animal remains, only Type BN-II increased reaching values similar to BN1. Charcoal slightly declined to intermediate values.

#### Zone BN6 (892 to 842 cm; 11,584-12,105 to 10,792-11,182 cal yr BP; Early Holocene)

Arboreal and nonarboreal pollen types returned to values similar to BN2 but this time the trees were represented by *Pinus* (up to 60%) and *Betula* (up to 20%). Nonarboreal elements did not show evident influx decreases and their decline in percentage was due to the influx increase of arboreal elements, which determined a significant increasing trend in PAR. Evergreen *Quercus* began to appear in a consistent fashion, and the first appearances of deciduous *Quercus* and *Corylus* were recorded at the end of the zone. Fungi did not show significant changes and algae declined again to values similar to BN2, in both percentage and influx, as well as in composition. Among animals remains, Cladocera showed relevant abundances for the first time. Charcoal significantly increased and peaked by the middle of the zone.

#### Zone BN7 (842 to 789 cm; 10,792-11,182 to 9578-10,362 cal yr BP; Early Holocene)

Zone characterized by the full dominance of arboreal pollen types (*Pinus*, *Betula*, *Corylus* and both *Quercus*) which reached values over 95%. Nonarboreal taxa were negligible, and only Poaceae attained values of ∼1%. This was due to a meaningful influx increase in tree elements, which resulted in maximum PAR values, and a slight influx decrease in nonarboreal taxa. All algae types declined to negligible values except *Cosmarium*, which slightly increased. Animal remains did not show significant changes, and *Pinus* stomata attained significant values throughout the zone. Charcoal influx remained high but highly fluctuating, and large particles (>150 μm) showed maximum values.

Individually, the first significant increases in some important elements, such as *Betula* (∼900 cm; 11.7-12.2 cal kyr BP), deciduous *Quercus* (∼850 cm; 11.0-11.2 cal kyr BP), *Corylus* (∼840 cm; 10.7-11.2 cal kyr BP) and *Ulmus* (∼820 cm; 10.4-11.0 cal kyr BP), are worth noting. Also important are the last significant appearances of relevant elements such as *Saxifraga* and Cichoridoideae (∼940 cm; 12.2-12.6 cal kyr BP), Chenopodiaceae (∼910 cm; 12.0-12.3 cal kyr BP), and *Artemisia* and Poaceae (∼890 cm; 11.5-12.1 cal kyr BP).

## 5. Discussion

This section is aimed at interpreting the above results in terms of vegetation dynamics and the potential influence of climatic drivers, notably temperature and moisture. The account is subdivided into two subsections: the first aims to identify the vegetation types that could have produced the pollen assemblages found, and the second aims to reconstruct the temporal trends of these vegetation types in relation to paleoclimatic shifts and fire.

### 5.1. Vegetation types

Modern pollen spectra around Bassa Nera are dominated by *Pinus* (∼80%) with small percentages of grasses (<10%), *Corylus* and deciduous *Quercus* (<1%) (Fig. 3). A similar assemblage has not been found in our pollen sequence, which indicates that the LGEH vegetation was significantly different from the present. This is not surprising, as previous glaciological studies demonstrated that the site was near the snowline during the Lateglacial [9]. However, no modern pollen analogs exist on the uppermost sites close to the present snowline (∼2800 m), which indicates that Lateglacial vegetation was also different from the present, even at the highest vegetation belts.

To unravel the PGEH vegetation types from Bassa Nera, a cluster analysis was performed on pollen taxa, avoiding scarce components with only sporadic occurrences. Four assemblages were identified, two representing open (nonforested) associations (O1 and O2) and two representing forests (F1 and F2) (Fig. 8). O1 was dominated by *Artemisia* and Poaceae, and O2 was dominated by *Saxifraga* and Cichoroideae. Forest associations showed patterns similar to the present: F1 was dominated by deciduous trees (notably *Betula* and *Corylus*) and coincided with Df, whereas F2 had a single component, *Pinus*, thus coinciding with Cf. The main difference with extant vegetation types was the Lateglacial abundance of O1 and O2 taxa, especially *Artemisia*, *Saxifraga* and Cichorioideae (but also Chenopodiaceae, other Asteraceae, Caryophyllaceae, *Ephedra*, Apiaceae and Plantago), which is unparalleled in modern pollen assemblages and deserves further consideration.

**Figure 8.**
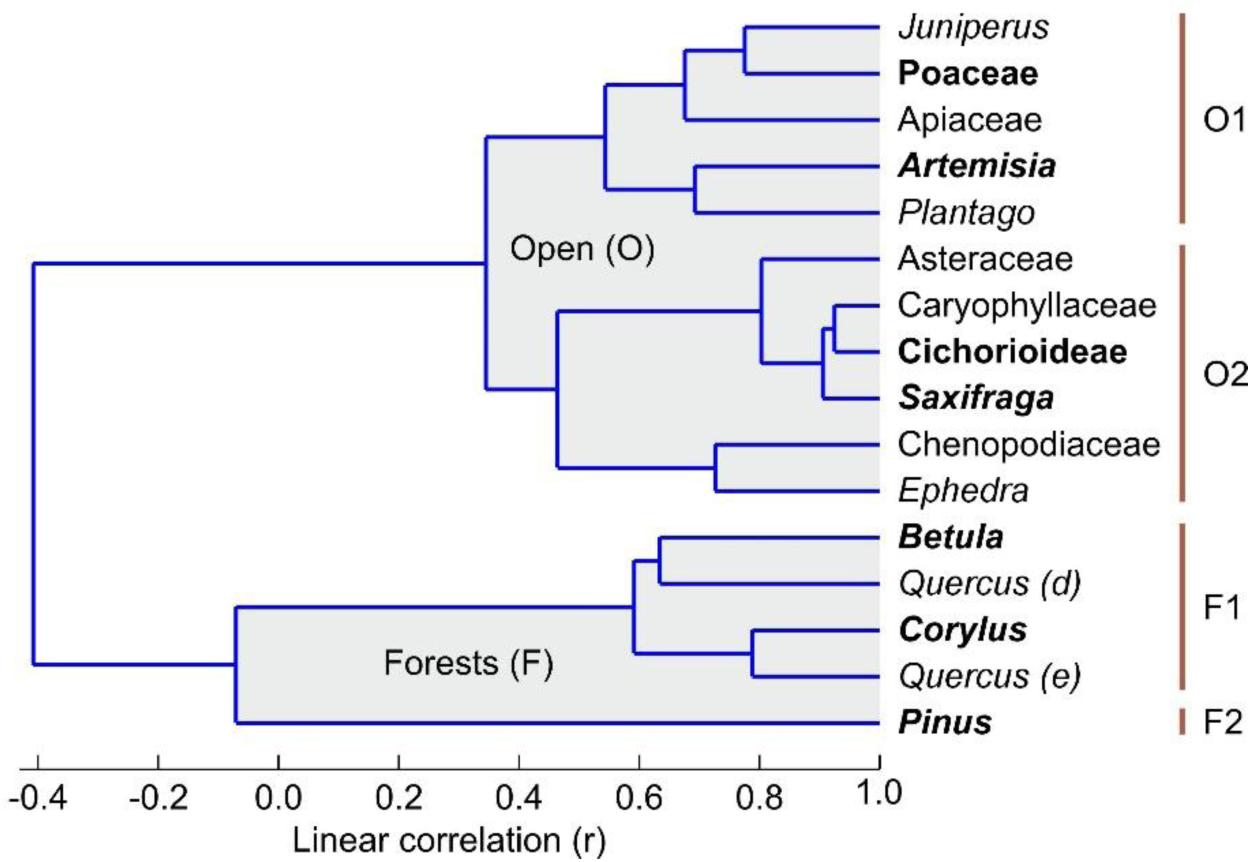
Cluster analysis using pollen taxa above 0.5% of the total. Dominant taxa of each association are in bold.

In the Mediterranean region, the indicator character of *Artemisia* pollen in past records has long been debated, which is linked to the difficulty of identifying the species involved using only pollen morphology. The pollen of this genus has traditionally been used to characterize glacial and other cold climatic phases, but according to Subally & Quétzel [54], the approximately 30 extant species of *Artemisia* living in the Mediterranean region bear a wide range of environmental conditions, and a strictly glacial character cannot be guaranteed, which is supported by increases in this pollen during interglacial and other warming phases. In addition, the concurrence of *Artemisia*, Chenopodiaceae and *Plantago* pollen has been considered strong evidence for regional grazing in the Pyrenees, whereas Cichorioideae has been linked to local grazing activities [55]. In our case, however, the absence of archeological and paleoecological evidence for human impact during the period of study, along with the lack of fungal spores from coprophilous fungi as indicators of grazing activities, strongly argues against this possibility in the Bassa Nera Lateglacial.

In the Pyrenees, Lateglacial pollen assemblages dominated by *Artemisia*, Poaceae and Chenopodiaceae have traditionally been interpreted in terms of cold and arid steppe-like vegetation on the basis of similarities with modern pollen spectra from this vegetation type in eastern Europe [10,15]. This interpretation was further extended to the adjacent Pyrenean lowlands [56] and the whole Iberian Peninsula [17,19], regardless of elevation. In some cases, however, steppe vegetation is not mentioned, and the association *Artemisia*-Chenopodiaceae (along with several Caryophyllaceae and Cichorioideae, among others) is considered to be characteristic of pioneering vegetation on open mineral soils, whereas Poaceae is considered to be dominant in closed herb vegetation [57]. Notably, these associations are similar to our O1 and O2 assemblages, but in our case, no modern analogs for these pollen assemblages have been found (Fig. 3). O1 differs from alpine meadow assemblages by the abundance of *Artemisia* and the presence of *Juniperus*, whereas O2 is lacking at all.

Assemblage O1 could represent herbaceous vegetation with *Artemisia* species characteristic of periglacial climates. Presently, two species of this genus, *A. eriantha* and *A. gabriellae*, occur at the highest elevations of the Pyrenean alpine belt; however, they are restricted to rocky habitats and do not occur on alpine meadows [58]. The shrubby *Juniperus* is represented in the alpine and subalpine stages by *J. communis* subsp. *nana* and *J. sabina*, respectively, both typical of relatively warm and humid environments. It seems unlikely that the same *Artemisia* and *Juniperus* species were present in extremely cold Lateglacial conditions. In addition to Poaceae, *Plantago*, represented by three species (*P. alpina*, *P. media* and *P. monosperma*), is the only component of this assemblage commonly growing on alpine meadows. Apiaceae pollen could not be identified, but it is mostly of the *Eryngium* type, which is represented in the high mountain meadows by *E. alpinum*, *E. bourgatii* and *E. campestre*. In summary, this pollen assemblage might be representative of uppermost alpine meadows with scattered shrubs with no modern analogs, probably of steppic nature due to the presence and abundance of unknown *Artemisia* and *Juniperus* species. With the available evidence, however, it is not possible to know whether this vegetation type is similar to the modern eastern European steppes because of the lack of sufficient taxonomic resolution (genera or families, at most). Therefore, this pollen assemblage will be considered to represent periglacial herbaceous vegetation with no further biogeographical considerations.

Assemblage O2 is even more difficult to interpret. One of the dominant taxa, *Saxifraga*, is represented in the Pyrenean highlands by numerous species, many of them living in rocky environments close to the snowline, situated at ∼2800 m elevation [58]. The codominant Cichorioideae could correspond to several genera with different ecological requirements, which is also true for other Asteraceae and Caryophyllaceae. The Chenopodiaceae are represented mainly by *Chenopodium bonus-henricus*, a nitrophilous megaphorb growing on forest clearings and ruderal sites. *Ephedra* is mostly represented by the *E. distachya* type, which is absent from the Pyrenean highlands and occurs mainly on southern dry lowland steppes. The lack of sufficient taxonomic resolution and the fact that a similar association has not been documented in extant plant communities or pollen assemblages [32] makes O2 difficult to interpret in terms of vegetation type. A number of O2 elements (notably Cichorioideae, Caryophyllaceae, Chenopodiaceae and *Ephedra*) have been considered in the Pyrenean and Iberian Lateglacial literature as characteristic of cold/dry Eurasian steppes and interpreted in combination with the O1 elements. However, the clear separation of O1 and O2 assemblages in our Bassa Nera record could have some paleoecological significance that remains unknown. Therefore, this pollen assemblage is considered here to represent an unknown high mountain plant association with no modern analogs.

### 5.2. Climate changes, vegetation shifts and fire

The relationship between the identified pollen assemblages and the independent paleoclimatic records is analyzed here using the abovementioned Ostolo and Seso speleothem records for temperature and moisture, respectively. The main departures of the Ostolo record with respect to the NGRIP2 standard were a small offset at the end of GS-2.1a (i.e., the Bølling warming commenced a couple of centuries before in OST2) and the absence of minor GI-1d, GI-1c2 and GI-1b coolings, along with the occurrence of a cooling event during the GI-1c1 Greenland warming (Fig. 1). The remaining trends are in good agreement with the ice core records, including the GS-1 (YD) and the 11.4 cold event. Regarding moisture, the GS-1/YD showed a conspicuous trend toward wet climates in the Seso record (Fig. 1), coinciding with another speleothem record from the French Moulis cave (Fig. 1) reporting humid conditions during the same period [47]. The occurrence of wet climates on Bassa Nera during the GS-1/YD is supported by the increase in freshwater algae influx, a pollen-independent proxy for water volume/level, which remained at lower values before and after this phase (Fig. 9). The significant increase in sediment accumulation rates occurred during the GS-1/YD (YDsa: Fig. 4) may also suggest the occurrence of local humid conditions. Given the good agreement between pollen zones and the INTIMATE event stratigraphy (Fig. 5), comparisons of the Bassa Nera palynological record with the available paleoenvironmental proxies will be conducted using the corresponding paleoclimatic phases (Fig. 9).

**Figure 9.**
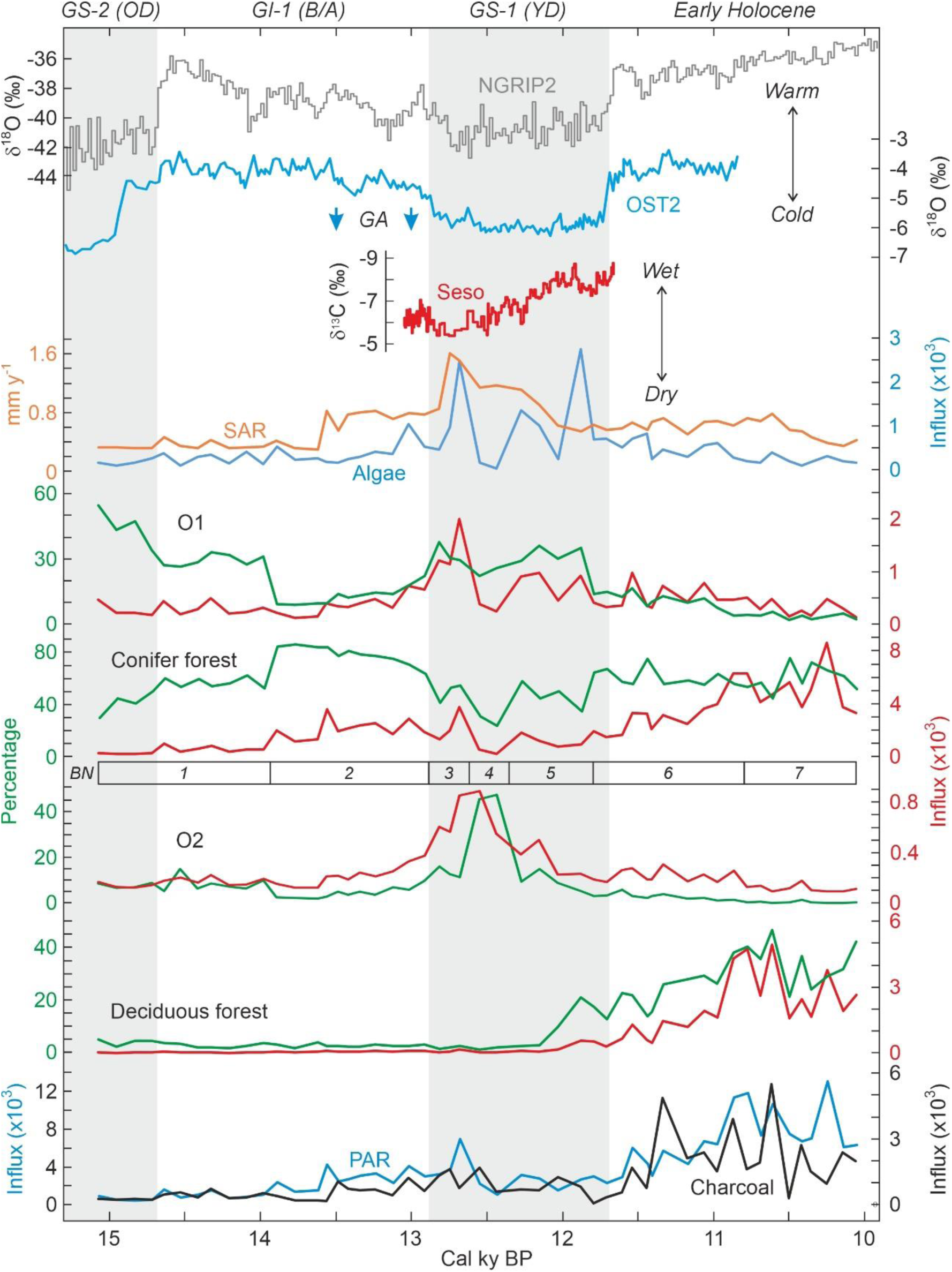
Comparison of pollen assemblages defined by cluster analysis (Fig. 8) with proxies for temperature, moisture availability and fire (Figs. 1 and 7). Pollen assemblages are presented in percentage and influx units (grains cm^-2^ y^-1^), sorted chronologically according to their respective maxima. BN, pollen zones; GA, glacier advances in the adjacent Ruda valley (Fig. 1); PAR, pollen accumulation rates; SAR, sediment accumulation rates.

**Figure 10.**
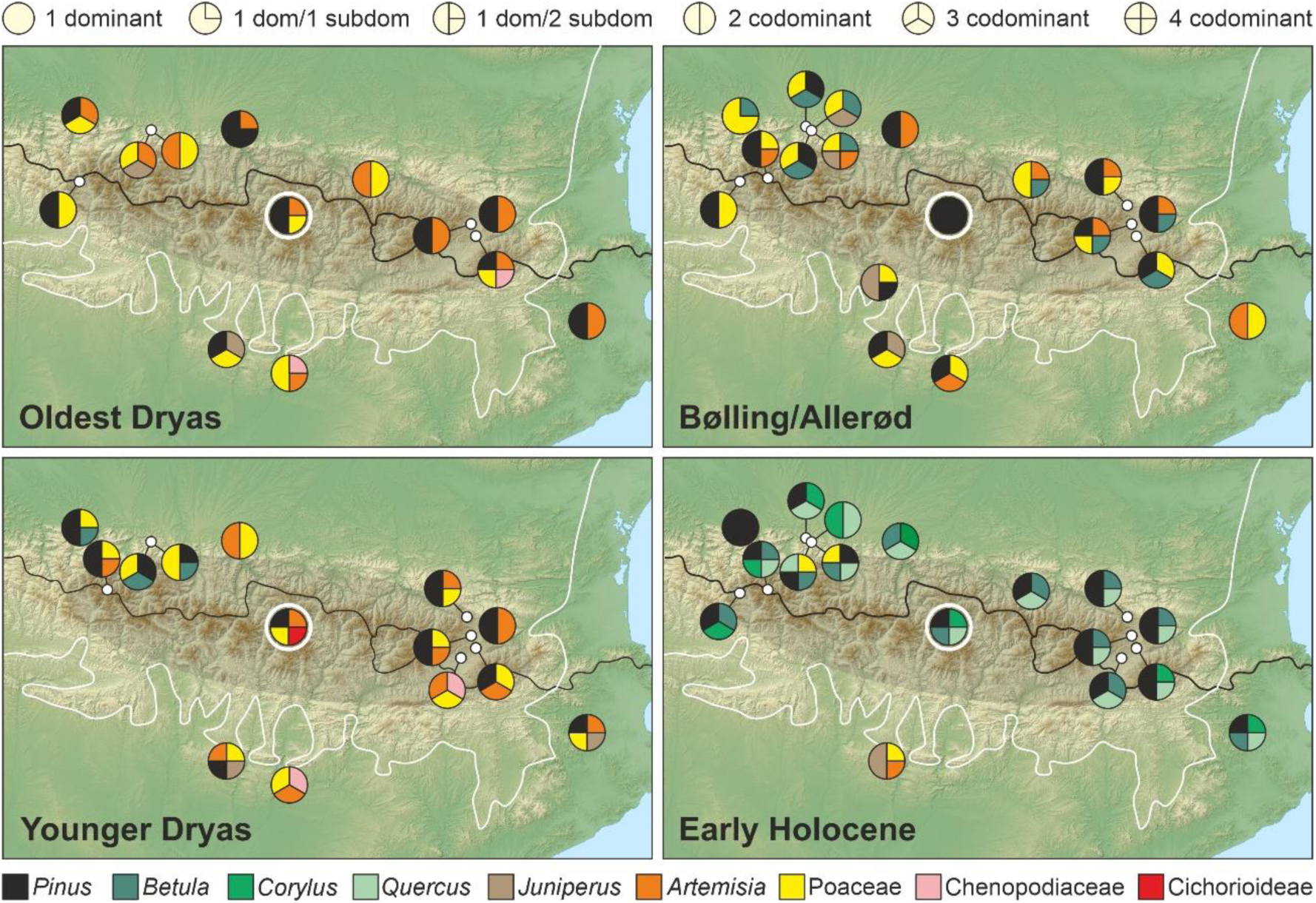
Sketch map with the main components of pollen assemblages from the LGEH stadials/interstadials in the different Pyrenean and lowland sites analyzed (see Fig. 2 and Appendix 1 for details). The Bassa Nera record is encircled by a white circle for easier location. As in Fig. 2, the black line is the border between France (northern Pyrenean slope) and Spain (southern Pyrenean slope), and the white line marks the boundary between Eurosiberian and Mediterranean biogeographical regions. Localities outside the shadow area are lowland records. Forest taxa are in black (Cf) and tones of green (Df and Sf) for better appreciation.

#### 5.2.1. GS-2 (Oldest Dryas)

Initial deglaciation of the Pla dera Muntanheta under still cold, although no longer glacial, climates and incipient sedimentation in a proglacial (seasonal?) pond in front of the receding moraines. Periglacial landscapes with dominant permafrost conditions with a poor plant cover of small scattered grassy *Artemisia* stands (O1) on suitable microenvironments. Pine forests were downslope, at elevations significantly lower than today. Indeed, the treeline was about 1500 m below the present, as suggested by *Pinus* pollen percentages (compare with Fig. 3), which would be compatible with average temperatures of the order of 6 °C or more below the present. By the time, deciduous forests were probably too far, probably in the basal lowlands, for their pollen to be wind-transported to high-mountain environments and the O2 vegetation had not yet colonized the site.

#### 5.2.2. GI-1 (Bølling/Allerød)

The abrupt global warming recorded by 14.7 cal kyr BP (∼15 cal kyr BP in the western Pyrenees) did not result in a significant landscape change on Bassa Nera, where O1 vegetation was declining and the downslope pine forests were rising, but only in relative abundance. The main vegetation shift took place about 0.5-1 cal kyr after temperature stabilization, in the middle of the interstadial (∼14 cal kyr BP, the boundary between pollen zones BN1 and BN2), when pine forests abruptly increased, although PAR values were still lower to support the presence of conifer forests on the coring site. Actually, pine forest percentages indicated a treeline situated approximately 800-1000 m below the present, which suggests a ∼500 upward shift with respect to the GS-2/Oldest Dryas stadial. After ∼14 cal kyr BP, temperatures started to gradually decrease, which was paralleled by a similar decline of conifer forests and rising O1 and O2 trends (Pollen Zone BN2) compatible with a local increase in plant cover on Bassa Nera. Notably, the two glacier readvances recorded in the adjacent Ruda valley at 13.5 and 13.0 cal kyr BP [9] occurred during this phase and approximately coincided with two significant accelerations of sediment accumulation rates (SAR), which would suggest enhanced glacial erosion (winter) and increased sediment transport of the eroded materials (summer) to the pond.

#### 5.2.3. GS-1 (Younger Dryas)

The onset of this stadial was marked by a distinct temperature decline and a major local runoff/moisture increase, as suggested by abrupt increments in SAR and algae influx. Temperatures remained low during the whole stadial but moisture experienced some short-lived (centennial) although significant changes. Indeed, algae decreased again by ∼12.6 cal kyr BP and peaked two more times by ∼12.3 and ∼11.9 cal kyr BP, which would indicate fluctuating water volumes/levels in the pond within a generally wetter climate. SAR showed a different pattern consisting of a continuous decline throughout the GS-1/YD, which would suggest a decreasing importance of winter erosion and/or summer melting, likely due to increased ice accumulation. This, together the rising moisture trend in the Seso record [8] and the occurrence of humid conditions, as deduced from enhanced speleothem growth, in the Moulis cave [47], in the southern and northern lowlands, respectively, strongly suggest cold and wet regional climates, although with eventual short-term local oscillations due to particular site features, during this stadial.

Moisture oscillations were paralleled by major vegetation changes. The first local moisture increase (∼12.7 cal kyr BP) coincided with an outstanding peak in O1 (Pollen Zone BN3) followed by a sudden decline coeval with a rapid increment of O2 (Pollen Zone BN4), coinciding with an equally rapid and intense moisture decline. This suggests that O1 would be favored by wet climates, whereas O2 would be more linked to dry conditions, at least under maintained cold climates. Subsequent moisture peaks had similar consequences, though less intense, for O1 (Pollen Zone BN5) but O2 never recovered after its BN4/BN5 decline. Therefore, hydroclimate would be more influencing than temperature in determining local vegetation types and cover. Indeed, temperature remained relatively stable during the whole stadial and affected mainly the regional vegetation, especially conifer forests, which were lowered to elevations similar to the GS-2 (OD) ones. Hydroclimate could also have had some influence on deciduous forests, which began to increase during the last moisture peak (Pollen Zone BN5) that occurred by 11.9 cal kyr BP, approximately two centuries before the abrupt warming that initiated the Holocene, when climates were regionally wet, as indicated by the Seso and Moulis speleothem records.

The occurrence of plant communities dominated by *Artemisia* and Poaceae (O1) under wet conditions contrasts with traditional interpretations of these associations as indicative of dry steppes, on the basis of biogeographical inferences [10]. On the one hand, the already discussed lack of sufficient taxonomic resolution for *Artemisia* pollen records, along with the wide arrangement of environments where the species of this genus may live, hinders specific paleoenvironmental inferences such as those attempted traditionally [54]. On the other hand, it should be noted that, in periglacial environments, wet climates and water stress can coexist, as moisture availability may be reduced by permanent freezing (permafrost) rather than by decreases in the hydrological balance characteristic of arid climates. Indeed, the vegetation of these environments, also known as periglacial deserts, occurs on highland tundra-like wet biomes [59,60]. Care must be taken when assigning well-defined vegetational terms to unknown past vegetation types. In the case of the Pyrenees, the use of the term “steppe” for the Glacial and Lateglacial open vegetation may implicitly imply the existence of specific climatic, ecological and biogeographical conditions that are not necessarily supported by the available paleoclimatic evidence. In the long run, these terms and the associated environmental features may be consolidated by repetition and become established as paradigms in the scientific literature. In this way, unwarranted paleoenvironmental reconstructions may become axiomatic and foster circular interpretations when analyzing the response of vegetation to LGEH climatic shifts.

#### 5.2.4. Early Holocene

The initial Holocene warming was characterized by the drastic reduction, and eventually the disappearance, of O1 and O2 vegetation types and the gradual increase of conifer and deciduous forests, which were likely ascending due to the warming. The situation of conifer forests was similar to the GI-2/BA interstadial, when the treeline was 800-1000 m below the present. The well-known high dispersion power of pine pollen prevents us from knowing whether conifer forests dominated by this tree grew on the coring site during the Early Holocene. Indeed, no significant differences were found in pine pollen percentages at elevations with conifer forests and above the treeline (Fig. 3). In this case, the presence of *Pinus* stomata can provide support, as they are deposited mostly locally and are morphologically distinct [61]. These stomata were absent until pollen zone BN7, except for a couple of single occurrences in BN4/BN5 (Fig. 6), which suggests that conifer forests were not present in the coring site until the Early Holocene, since ∼11 cal yr BP onward. High PAR values also support this interpretation.

The case of deciduous forests was very different, as they were absent in the GI-2/BA stadial but very well developed in the Early Holocene. Actually, the pollen signal of these Early Holocene deciduous forests was similar to today’s montane stage (Fig. 3), situated 500-1000 m below under average temperatures 3-6 °C higher than in Bassa Nera. Although such a general elevational shift is hard to accept, it is difficult to escape the idea that deciduous forests were closer to Bassa Nera than they are today. A possible explanation is that these forests expanded from nearby small refugia with suitable microclimatic conditions – i.e., microrefugia [62] – as formerly proposed for the northern Pyrenean slope [10,11]. A number of these microrefugia for forest trees within Pleistocene glacial and periglacial environments have been documented in SW France using edaphic charcoal analysis [63]. In the case of evergreen *Quercus*, this explanation is unlikely, as its climatic habitat lies ∼2000 m below the snowline. However, the upward dispersion of its pollen is noteworthy, as demonstrated by the modern-analog study in the Aiguamog valley, where evergreen oaks are absent but their pollen occurs along the whole transect up to 2600 m elevation [32]. It should be clarified that the occurrence of evergreen *Quercus* within the Df assemblage is not anomalous, as mixed deciduous and evergreen oak forests are common in the montane stage, which is reflected in modern pollen assemblages [32].

#### 5.2.5. Fire incidence

Fires began to be important in the Early Holocene, as indicated by the charcoal record. Interestingly, fire incidence (charcoal) and total vegetation cover (PAR) showed a statistically significant correlation (Table 2) for the whole Bassa Nera record, which suggests that fuel availability (in the form of biomass accumulation), along with higher temperatures, was a major driver for fire, as previously noted in other localities from the northern and southern Pyrenean sides [14,20]. It should be noted that the main bulk of charcoal particles found in this study were below the boundary for regional vs. local fire events (150 μm), and therefore, they record mostly regional fires, likely by burning of lower elevation forests. Charcoal particles larger than 150 μm were comparatively scarce and only increased in zone BN7, coinciding with the appearance of *Pinus* stomata (Fig. 6), which suggests that the burning of in situ pine forests did not occur before those dates. Only forests, especially deciduous forests, showed significant correlations with charcoal (Table 2), which suggests that they provided the main fuel for regional forest fires.

**Table 2.** Pearson correlation coefficients between the pollen assemblages defined by cluster analysis (Fig. 8) and charcoal using influx units (grains/particles cm^-2^ y^-1^). Significant correlations in bold.

|  | **Charcoal** | **p (uncorr)** |
| --- | --- | --- |
| PAR | **0.6249** | 9.5 10<sup>-7</sup> |
| Conifer forest | **0.4845** | 0.0003 |
| Deciduous forest | **0.6999** | 1.1 10<sup>-8</sup> |
| O1 | 0.1453 | 0.3088 |
| O2 | 0.0162 | 0.9103 |

## 6. Regional comparisons

Pyrenean-wide comparisons among pollen assemblages were carried out for each LGEH stadial/intestadial considering general climatic (wet Atlantic vs. dry Mediterranean), biogeographical (Eurosiberian vs. Mediterranean) and elevational (highlands vs. lowland) gradients (Fig. 9). Independent paleoclimatic reconstructions are scarce in surveyed localities, and most paleoenvironmental inferences are based on the same palynological proxies used for vegetation reconstruction, which hinders detailed comparisons of vegetation responses to past climatic changes. To homogenize nomenclature, we used the classical terminology (Oldest Dryas, Bølling/Allerød, Younger Dryas), as the updated terminology (GS-2, GI-1, GS-1) was unavailable in former studies. Sites with dating problems and/or poorly defined LGEH phases (Appendix 1) were not considered in this comparison.

### 6.1. Oldest Dryas

During this stadial, our record (BN) shows an intermediate situation between the Atlantic (western) and Mediterranean (eastern) sides. Indeed, open BN landscapes were codominated by Poaceae and *Artemisia*, the first being more abundant in the western open formations and the second being more important in nonforested areas from the eastern sector. Forest communities did not show significant differences and were dominated by *Pinus* on both sides. In the lowlands, the situation was also intermediate but open landscapes predominate over forested formations, except for a northern locality (Br), where *Pinus* was dominant. Sites from the Mediterranean biome did not show significant departures from Eurosiberian localities and were more consistent with the east‒west gradient. The general impression is of relatively open landscapes dominated by Poaceae and *Artemisia* – with *Juniperus* in the west and Chenopodiaceae in the east – intermingled with conifer forests everywhere.

### 6.2. Bølling/Allerød

During this interstadial, BN was fully dominated by pine pollen, which strongly contrasts with eastern-western Pyrenean sides and with lowlands, which were covered by mosaic vegetation. In general, the former Poaceae-*Artemisia* dipole was less pronounced and the forest cover increased, but at that time *Betula* was as important as *Pinus* everywhere, except in the southern and eastern lowlands, where *Betula* was not an important element of the vegetation. Also noteworthy is the increase in *Juniperus* in the northern and southern lowlands from the central and western sectors. In general, during the B/A, major differences were recorded between the northern and southern Pyrenean sides, rather than on the east‒west, highland-lowland or Eurosiberian-Mediterranean gradients.

### 6.3. Younger Dryas

During this stadial, open vegetation became more dominant, and the east‒west gradient accentuated due to the reinforcement of the Poaceae-*Artemisia* dipole (which was even more pronounced than in the OD) and to the newly emerged contrast between western and eastern forests due to the importance of *Betula* in the former and its absence in the latter. In this case, the BN record was intermediate regarding open formations but more similar to the eastern sector because of the absence of *Betula* as an important component. In addition, our record differed from all other Pyrenean localities in the presence of the Sc (*Saxifraga*-Cichorioideae) assemblage. No evident differences between highlands and lowlands or between sites from Eurosiberian and Mediterranean regions were observed.

### 6.4. Early Holocene

A major regional vegetation shift took place at the onset of the Holocene due to the full dominance of forest formations, except for a southern locality (Es), where stadial-like open landscapes remained. This general forest increase was due to the influx of *Betula*, *Corylus* and *Quercus* rather than to an expansion of conifer forests. In this case, BN experienced the same general replacement of open vegetation by newly arrived forests, which showed similar patterns across the east‒west, highland-lowland and Eurosiberian-Mediterranean gradients, except for a few western lowland sites where grasslands were still important. Some minor differences between western and eastern forests were the importance of *Corylus* in the former and *Betula* in the latter, with BN in an intermediate position.

## 7. Final remarks and future prospects

In addition to providing the first continuous Lateglacial-Early Holocene palynological record for the Iberian Pyrenees at centennial resolution, this study emphasizes the need for improving paleoclimatic interpretations, especially in reference to the use of pollen-independent proxies to avoid circularity. In particular, inferences about the occurrence of cold/dry climates based on the occurrence of *Artemisia*-Poaceae steppes during the Younger Dryas are not consistent with independent paleoclimatic evidence suggesting the occurrence of wet climates for this stadial. Unfortunately, the lack of pollen-independent paleoclimatic records is insufficient for a sound assessment at regional level. Emphasis should be placed on the development of regional paleoclimatic studies based on suitable physico-chemical and biological proxies before analyzing the response of vegetation to paleoenvironmental changes, rather than inferring paleoclimates from pollen reconstructions, as traditionally attempted. This work has also suggested the potential occurrence of climatic oscillations during the GS-1/YD stadial that are worth to be studied in more detail at higher resolution (decadal or less). In addition, independent paleohydrological reconstructions for Lateglacial phases other than the GS-1/YD are lacking in the Pyrenees and should be encouraged for a more consistent regional paleoclimatic view.

Finally, it would be interesting to extend regional paleoclimatic and paleoecological comparisons to the continental ambit for a better appraisal of the influence of climatic, biogeographical and anthropogenic (when present) factors on the composition and temporal trends of LGEH montane vegetation across Europe. The use of the European Pollen Database (https://epdweblog.org/) would be especially useful in this sense but should be complemented with more regional compilations similar to those existing for the Iberian Peninsula [64], which have been demonstrated to be much more exhaustive and useful than the available global databases for this type of reconstruction [65].

## Data Availability Statement

Raw data available upon request to V.R. or T.V.V.

## Acknowledgments

Fieldwork collaborators, Encarni Montoya, Nick Loughlin, Arantza Lara, Núria Cañellas-Boltà, Sandra Garcés-Pastor and Julià López-Vila; modern climatic data, Javier Sigró. Special thanks to Génesis Hernández for the high-quality sample processing, which greatly facilitated palynological work.

---

**Appendix 1.**
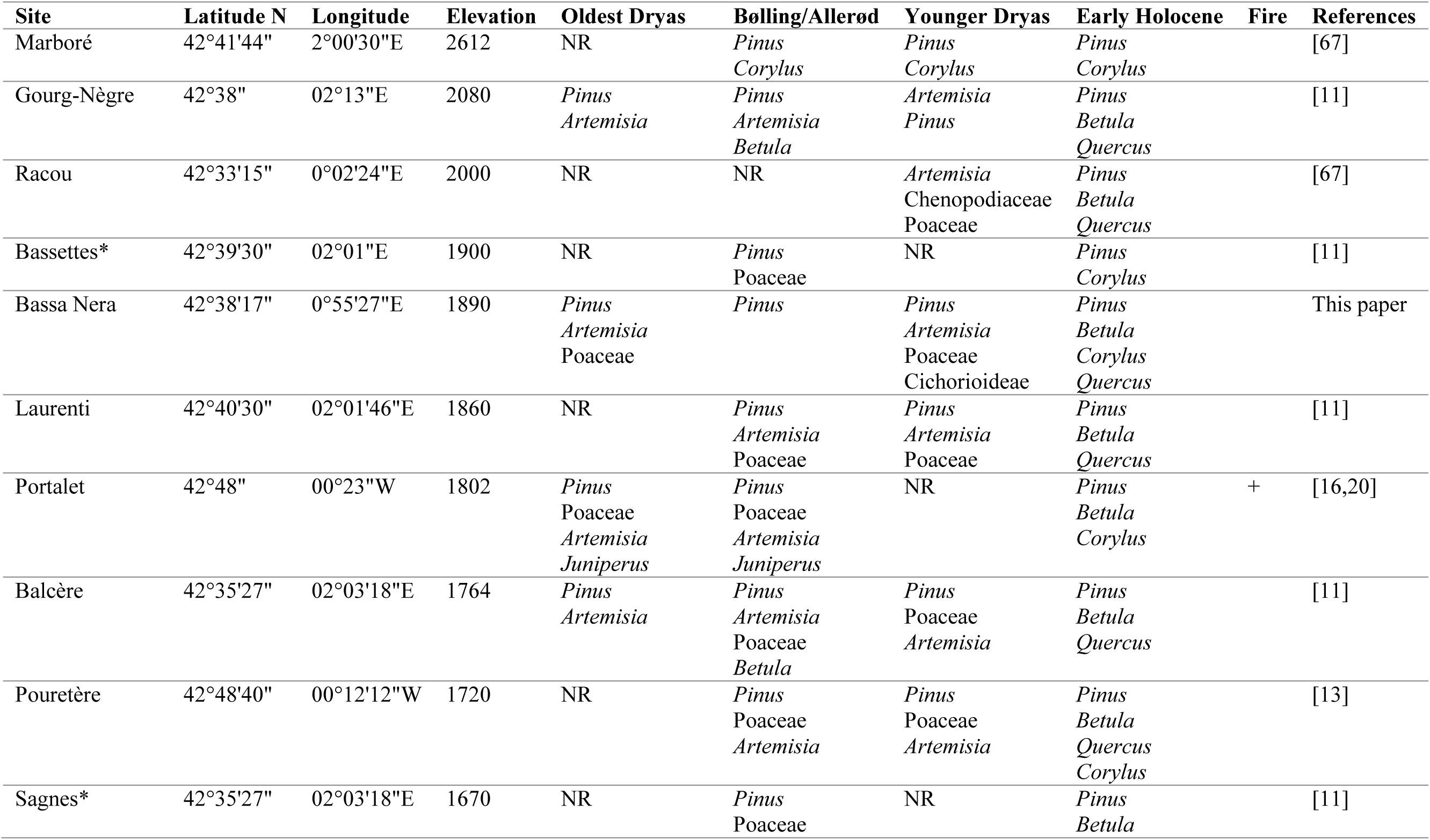
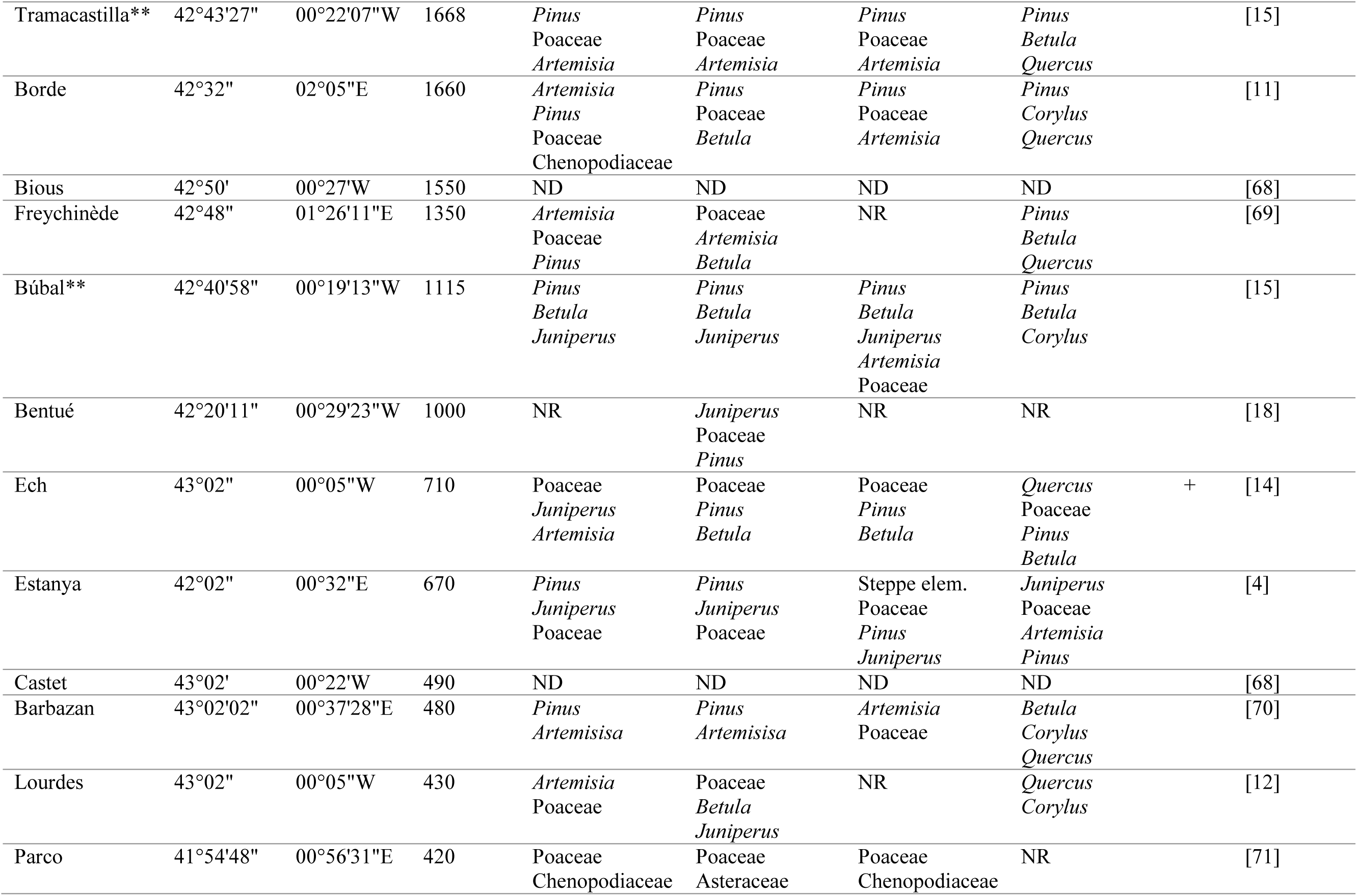
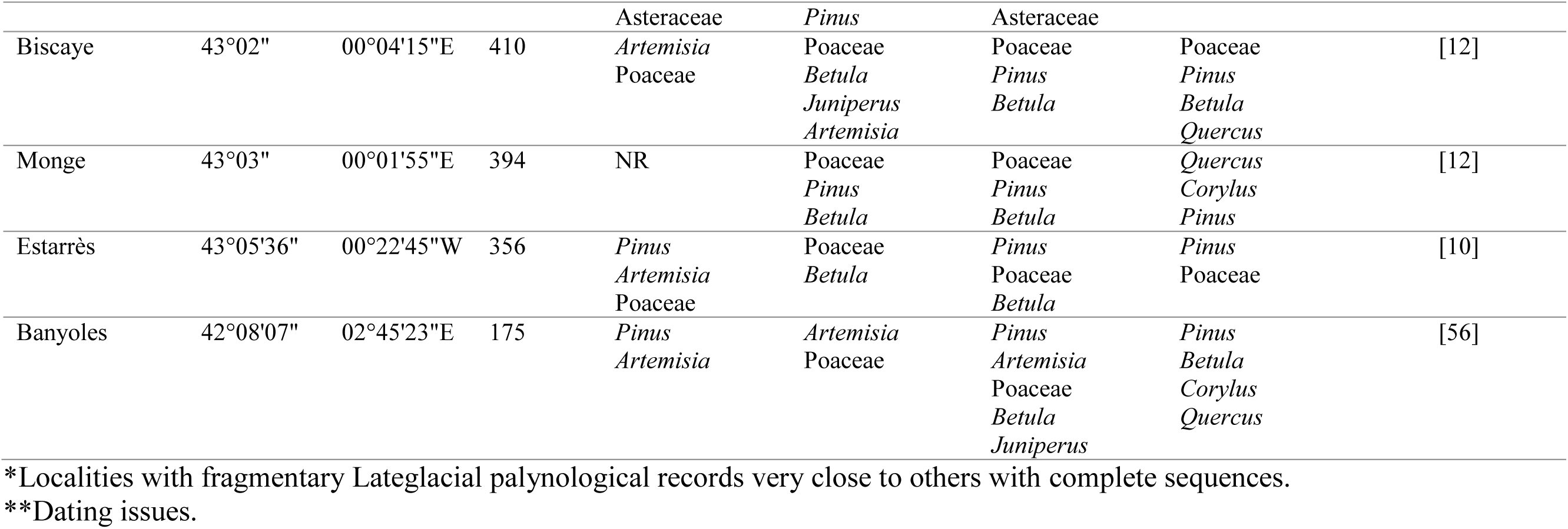
Details of the Lateglacial coring localities with published palynological records, sorted by elevation, including the major pollen taxa used to assemble Fig.listed in top-down dominance order. Palynological information has been taken directly from the pollen diagrams displayed in the cited references. NR, not recorded; ND, possibly present but not defined.

